# Identification of QTLs for high grain yield and component traits in New Plant Types of rice

**DOI:** 10.1101/2020.01.07.897330

**Authors:** Ravindra Donde, S. Mohapatra, S. Y. Baksh, B. Padhy, M. Mukherjee, S. Roy, K. Chattopadhyay, A. Anandan, P. Swain, K. K. Sahoo, O. N. Singh, L. Behera, S. K. Dash

## Abstract

A panel of 60 genotypes consisting of New Plant Types (NPTs) along with *indica*, *tropical* and *temperate japonica* genotypes were phenotypically evaluated for four seasons in irrigated situation for grain yield *per se* and component traits. Twenty NPT genotypes were found to be promising with an average grain yield of 5.45 to 8.8 t/ha. A total of 85 SSR markers were used in the study to identify QTLs associated with grain yield *per se* and related traits. Sixty-six (77.65%) markers were found to be polymorphic. The PIC values varied from 0.516 to 0.92 with an average of 0.704. A moderate level of genetic diversity (0.39) was detected among genotypes. Variation to the tune of 8% within genotypes, 68% among the genotypes within the population and 24% among the populations were observed (AMOVA). The association analysis using GLM and MLM models led to the identification of 30 and 10 SSR markers were associated with 70 and 16 QTLs, respectively. Thirty novel QTLs linked with 16 SSRs were identified to be associated with eleven traits, namely, tiller number (*qTL-6.1, qTL-11.1, qTL-4.1*), panicle length (*qPL-1.1, qPL-5.1*, *qPL-7.1, qPL-8.1*), flag leaf length (*qFLL-8.1, qFLL-9.1*), flag leaf width (*qFLW-6.2, qFLW-5.1*, *qFLW-8.1, qFLW-7.1*), total no. of grains (*qTG-2.2, qTG-a7.1*), thousand-grain weight (*qTGW-a1.1, qTGW-a9.2, qTGW-5.1, qTGW-8.1*), fertile grains (*qFG-7.1*), seed length-breadth ratio (*qSlb-3.1*), plant height (*qPHT-6.1, qPHT-9.1)*, days to 50% flowering (*qFD-1.1*) and grain yield per se (*qYLD-5.1, qYLD-6.1a, qYLD-11.1*). This information could be useful for identification of highly potential parents for development of transgressive segregants. Moreover, super rice genotypes could be developed through pyramiding of these QTLS for important yield traits for prospective increment in yield potentiality and breaking yield ceiling.

## Introduction

Rice (*Oryza sativa* L.) is a staple crop for more than 3.5 billion people in the globe. In current scenario, rice productivity is increasing rate at 1% per year which is less than the 2.4% per year rate required to double the global production by 2050 [1]. Considering a glimpse of the history, a quantum jump in productivity was achieved due to the green revolution in mid-sixties, which drastically enhanced the rice production of the world. However, a ceiling of productivity potentiality is reported by and large in semi-dwarf inbred *indica* genotypes since release of IR-8 [2], in spite of substantial improvement in yield stability, per day productivity and grain quality [3]. A breakthrough in productivity barrier is necessitated because of increasing competition for natural resources viz., land, water and others given population explosion coupled with expanding industrialization, urbanization and diversion of agricultural land [1,4,5]. This is still aggravated with the abnormal change in weather and climate with significant influence on crop productivity and quality [6, 7].

Rice scientists are facing many challenges for doubling rice production by 2050. Irrigated rice has a share of 75% of total rice production in the world, although it has a share of about 55% of the total rice area [2]. Therefore, improvements and modification in this ecology are supposed to have significant impact on rice productivity in future. During the past decade, there has been a significant slowdown in the production potential of modern rice cultivars. In this context, physiologists and breeder hypothesised that this stagnation could be overcome by improving the plant type. The existing plant type bears several unproductive tillers in high tillering type and limited sink size i.e., small panicles. The excessive leaf area causes mutual shading, low light and a reduction in canopy photosynthesis [8, 9]. Apart from that, there are several bottlenecks viz., spikelet sterility, short panicle length, limited grain numbers, lodging susceptibility, etc. Moreover, there is also the loss of genetic diversity in improved varieties for which breeders are facing difficulties in finding divergent gene pools. Modern high yielding rice varieties have been associated with some unfavourable traits/alleles, which may be sensitive to biotic and abiotic stresses and may be responsible for lowering grain yield [9].

In this context, IRRI scientists developed “New Plant Type” (NPT-2^nd^ generation) by recombining some suitable features of *tropical japonicas* with *indica* [9]. The main idea behind of 2^nd^ generation NPT was development of high yielding super rice varieties, which can able to produce significant-high yield along with stability. Some of the NPTs performed exceedingly well and produced even more than 10t ha^−1^ at Philippines [2]. During the process of development, some of advanced generation segregating materials were shared with NRRI, India. The materials were subjected to further selection at NRRI for irrigated ecology as appreciable variability was still available, with an objective of development of promising NPTs suitable for the climate specific to eastern region in particular and country in general. Trait specific selections were exercised for few generations to establish fixed lines, i.e., NPT selections (NPTs). In this context, NPTs were evaluated systematically under observational yield trial (OYT) for one season and the number was narrowed down. It was followed by Advanced Yield Trial (AYT) for four wet seasons at NRRI. Some of the highly promising NPTs were identified with good agronomic traits like higher grain number per panicle, panicle length, panicle weight, grain size, ear bearing tiller number along with ideal plant height. Some of NPTs performed exceptionally well and showed the productivity of more than 10.0 t ha^−1^ during dry season 2011 [10].

With this backdrop, we wish to proceed for development of still higher yielding genotypes or super rice kind of crop ideotype utilizing the existing set of highly promising NPTs, which should have the productivity potential, at least 20% higher than the popular rice and check varieties. The target was utilization of one of the most promising gene pools through conventional as well as molecular approach. The focus is to accumulate the thousands of minor QTLs with additive genetic variance along with major ones. Here, the extent of genetic variation and relationships between genotypes are more important for designing effective breeding strategy [11].

The association mapping (AM) being useful tools in identifying QTLs/genes associated with different traits in plant species. It utilizes natural variation [12], hence supposed to have great potential to evaluate and characterize a wide range of alleles. Several researchers have been reported the utility of association analysis in the identification of QTLs for different traits in rice, viz., Grain yield [13], grain yield under water deficit [14], deep root mass and the number of deep roots [15], grain quality traits [16], agronomic traits [17, 18], grain yield under reproductive drought stress [19], panicle architecture and spikelet’s/ panicle [20], plant height and grain yield [21].

There are not sufficient reports on NPT for QTLs association on grain yield and yield-related complex traits. The genes/ QTLs related to high grain yield would be of great help in breaking yield ceiling. Moreover, it would be beneficial in identifying traits specific donors for designing effective breeding strategy for the development of super rice. The present study was undertaken to identify QTLs associate with grain yield and yield-related agronomic traits using diverse genotypes.

## Materials and Methods

### Plant Materials

Sixty rice genotypes panel, including 48 NPTs, six *indica* varieties, and significantly diverse, distinct three temperate *japonica* and three tropical *japonica* varieties were used for identification of QTLs for 11 yield-related traits through association studies (S1 Table). *Indica* varieties are highly popular, well adapted, released and notified for cultivation in different states of India. A total of 41 NPT populations were collected from IRRI at the advance segregating stage. From those populations, conscious trait specific single plant selections (SPS) were made basically for yield-related traits for few generations and finally ∼500 promising fixed SPS were identified and evaluated in OYT at ICAR-National Rice Research Institute (NRRI) (coordinates 20.4539° N, 85.9349° E). Subsequently, the number was drastically narrowed down to 48 strictly basing on yield and important agronomic traits. Forty-eight best performing NPTs have been evaluated in four environments along with 12 checks (6 *indica*, 3 *tropical japonica* and 3 *temperate japonica*) and these were further studied for molecular diversity and QTL association.

### Phenotyping

All the 60 genotypes were grown in two replications following Randomized Complete Block Design (RCBD) during wet seasons of 2011, 2012, 2013 and 2014 (S2 Table). The phenotypic data of yield *per se* and yield-related traits were recorded at different phenological stages. Normal management practices and pest protection measures were followed during crop growth. The genotypes were harvested separately after 30 to 35 days of flowering. The post-harvest data were recorded after the crops being harvested, then threshed and dried under sunlight. This study primarily focused on 11 yield and yield-related traits, namely, days to 50% flowering (DFF), plant height (PH), tiller number (TL), panicle length (PL), flag leaf length (FLL), flag leaf width (FLW), no of fertile grains (FG), total no of spikelets per panicle (TG), 1000-grain weight (TGW), seed length-breadth ratio (SLBR) and grain yield t/ha (YLD). The yield *per se* was measured by weighing the plot yield (4m^2^ each) at 13% moisture level and converted it to tons/ha. Other yield contributing traits were measured using standard procedure. The seed length-breadth ratio was measured using Anndarpan machine and software developed by CDAC, Govt. of India [22]. The phenotypic data were used for statistical analysis, namely, SD, CV, ANOVA, correlation, regression, and principal component analysis (PCA) Bi-plots using XLSTAT software (Addinsoft, Paris, France). The ClustVis, an online web tool (http://biit.cs.ut.ee/clustvis/) was used for analysis of phenotypic traits. The visualizing clustering of multivariate data of yield *per se* and yield-related traits were analyzed by Heat-map and PCA [23]. The ClustVisis mainly wrote in the Shiny web application framework by using several R package version 0.10.2.1 for R statistics software [16, 24].

### Genotyping

The genomic DNA was isolated from 3-4 gm of fresh leaf tissues of each rice genotype following Cetyl Trimethyl Ammonium Bromide (CTAB) method (Murray and Thompson, 1980). The extracted genomic DNA samples were dissolved in TE buffer (10 mM Tris-base, 1 mM EDTA, pH-8.0). The quality and quantity of DNA of each sample were checked by agarose gel electrophoresis and spectrophotometer. The SSR markers were selected on the basis of the previous report associated with different yield QTLs [25–31] and polymorphic contents (http://www.gramene.org). The polymerase chain reaction (PCR) was performed in a 20µl reaction mixture containing 5 pM (pico-mole) of forward and reverse primers of each SSR locus, 200 mM of each dNTP, 0.5 U of *Taq* DNA polymerase, 10 mM Tris-HCl (pH=8.3), 50 mM KCl and 1.5 mM MgCl_2_. The PCR amplification was carried out in a thermal cycler (Veriti 96, Applied Biosystems, USA) as per the following cycling parameters: initial denaturation at 94^0^ C for 3 min followed by 35 cycles of denaturation at 94^0^ C for 1 min, annealing at 55-67^0^ C (depending upon primer) for 1 min and extension at 72^0^ C for 1.5 min and final extension at 72^0^C for 5 min. The amplified products were separated on 2.5% – 3% agarose gels using 1x TBE buffer and stained with ethidium bromide (0.5 µg/µl). The gels were visualized under UV radiation and were photographed using a gel documentation system (G-Box, Syngene, USA) to detect amplified fragments. The size of amplified bands was determined based on the migration relative to molecular weight size markers (50 bp DNA ladder, MBI Fermentas, Lithuania).

#### Genetic Diversity

The amplified bands were scored as present (1) or absent (0) for each genotype and microsatellite marker combination. Each band was considered as an allele. The data were entered into a binary matrix as discrete variables and subsequently used for assessing allelic and molecular diversity such as number of total alleles (TA), unique alleles (UA), rare alleles (RA), expected alleles (Ne), polymorphism information content (PIC), gene diversity, homozygosity (Ho) and heterozygosity (He) by using Power-Marker Ver 3.25 [32]. The polymorphism information content (PIC) was calculated using the formula,“ 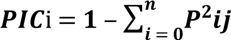” where Pij is the frequency of j^th^ allele for the i^th^ [33].

The genotypic data of 66 polymorphic markers were used for genetic diversity analysis. Jaccard’s similarity coefficient was calculated by using the NTSYS-PC software package [34, 35]. Cluster analysis was performed using UPGMA and sequential agglomerative hierarchal nested (SHAN) module of NTSYS-PC. The Nei’s pairwise genetic distance neighbour-joining [36] and Shannon’s diversity index (*I*) was calculated using POPGENE v 1.32 (http://www.ualberta.ca/fyeh) and MEGA 6 software. The Power-Marker was used for better visualization and understanding the clustering pattern of genotypes. The estimation of population differentiation among and within the genotypes was analyzed by Principal coordinates analysis (PCoA) and AMOVA by using software GeneAlEx 6 version 6.501 [37]. AMOVA was used to assess molecular variance within and between populations at 999 permutations.

#### STRUCTURE Analysis

Bayesian model-based clustering analysis available in STRUCTURE software 2.3.4 was used for data analysis to obtain possible population structure [38, 39]. The software provides the likelihood, classifies according to their population types, and assumes as K. The highest likelihood can be interpreted by the corresponding estimate of the basic number of clusters [39]. Each genotype was burned 10,000 and 150,000 steps followed by 100,000 and 150,000, respectively, using Monte Carlo Markov Chain replicates (MCMC). The K-value was run for 10 times with a K-value ranging from 1 to 10. The optimum K-value was determined by plotting the log posterior probability data to the given K-value. The ΔK value was estimated using the parameters described by Evanno et al.(2005) [40] using online software program Structure Harvester v6.0 (http://btismysore.in/strplot). In structure, the value of K is not constant because the distribution of L (K) does not show a clear cutoff point for the true K. An *ad hoc* measure of ΔK was used to detect the numbers of the subgroups. Some independent replicates, the admixture model and allele frequency model (length of burn-in + length of an MCMC repetitions x, number of independent replicates) were also calculated [13,39,41].

#### QTLs Association

The GLM, MLM, Quantile-Quantile (Q-Q) plot and Manhattan plot were used for association analysis of 11-grain yield and related traits by incorporating Q+K matrices using TASSEL version 5.2.9 [42]. The p-values at <0.005 level of significance were used to determine the significant association of SSR markers. In GLM and MLM, association analysis was performed at 1000 permutations for the correction of multiple testing [43, 44]. The False Discovery Rate (FDR) was calculated using SPSS statistical v20. (http://www-01.ibm.com/support/docview.wss?uid=swg21476447) at the 5% threshold level for multiple testing to standardise p-value [45]. The false-positive markers-traits association was controlled by applying models Q, K, and Q+K that were compared with each other using quantile-quantile (Q-Q) plot [46].

#### In-silico Study

The in-silico study was carried out for analysis of previously reported QTLs and genes associated with respective traits using computer and web-based servers. For this study, several web-servers were used i.e. http://www.gramene.org/, https://www.ncbi.nlm.nih.gov/ and https://rapdb.dna.affrc.go.jp/ etc. This study helps confirmation of QTLs and genes were associated in our rice population.

## Results

### Phenotypic Variation

The grain yield in rice is considered to be the most important trait in crop improvement. It controlled by several complex traits. The present study was a focus on ten yield-related traits that directly or indirectly control the grain yield. The set of 60 genotypes were phenotypically evaluated for grain yield and associated traits in irrigated situation in four consecutive wet seasons. A wide range of phenotypic variation was observed in all the grain yield and 10 yield-related traits (Table 1).

**Table 1.** The performance of 60 genotypes in four seasons based on grain yield and yield-related traits.

| Traits | Wet Seasons | Mean ± SD | CV (%) | Range | Correlations with yield | P-value |
| --- | --- | --- | --- | --- | --- | --- |
| DFF | 2011 | 100.99±8.61 | 8.53 | 72.5-126.0 | 0.34 | 0.006 |
| | 2012 | 101.78±8.02 | 7.88 | 75.78-123.74 | 0.49 | 0.000 |
| | 2013 | 104.44±6.09 | 5.83 | 80.5-116.5 | 0.35 | 0.005 |
|  | **2014** | 102.03±7.72 | 7.56 | 78.25-124.65 | **0.50** | **0.0001** |
|  | **Mean** | 102.31±7.13 | 6.97 | 76.76-121.47 | **0.48** | **0.000** |
| **PH** | **2011** | 109.38±18.24 | 16.67 | 60.65-181.9 | 0.38 | **0.001** |
|  | **2012** | 115.40±16.66 | 14.43 | 58.34-186.41 | **0.51** | **< 0.0001** |
|  | **2013** | 109.04±14.00 | 12.84 | 66.96-178.76 | **0.37** | **0.004** |
|  | **2014** | 107.92±13.84 | 12.82 | 76.55-186.93 | 0.18 | 0.1760 |
|  | **Mean** | 110.43±14.25 | 12.91 | 65.63-183.5 | **0.42** | **0.001** |
| **TL** | **2011** | 8.03±0.85 | 10.55 | 6.2-10.5 | 0.36 | **0.001** |
|  | **2012** | 8.71±1.40 | 16.13 | 5.83-12 | **0.31** | **0.015** |
|  | **2013** | 8.70±1.49 | 17.08 | 5-12.6 | **0.34** | **0.008** |
|  | **2014** | 7.43±0.85 | 11.41 | 5.625-9.55 | **0.44** | **0.0005** |
|  | **Mean** | 8.22±0.80 | 9.75 | 5.92-9.73 | **0.56** | **< 0.0001** |
| **PL** | **2011** | 28.21±3.12 | 11.08 | 18.25-33.15 | 0.68 | **< 0.0001** |
|  | **2012** | 28.24±3.27 | 11.57 | 17.19-34.8 | **0.67** | **< 0.0001** |
|  | **2013** | 27.60±2.74 | 9.94 | 21.27-32.65 | **0.51** | **< 0.0001** |
|  | **2014** | 27.12±2.00 | 7.37 | 21.26-31.6 | **0.41** | **0.0013** |
|  | **Mean** | 27.79±2.34 | 8.42 | 20.17-31.45 | **0.73** | **< 0.0001** |
| **FLL** | **2011** | 35.98±6.49 | 18.05 | 17.92-50.55 | 0.54 | **< 0.0001** |
|  | **2012** | 35.84±5.40 | 15.07 | 17.72-51.84 | **0.49** | **< 0.0001** |
|  | **2013** | 34.39±4.35 | 12.64 | 23.63-44.30 | **0.37** | **0.003** |
|  | **2014** | 34.64±3.72 | 10.75 | 26.51-46.12 | **0.46** | **0.0002** |
|  | **Mean** | **35.22±4.30** | 12.21 | 21.52-48.08 | 0.17 | 0.195 |
| **FLW** | **2011** | 1.43±0.23 | 16.35 | 0.65-1.81 | 0.53 | **< 0.0001** |
|  | **2012** | 1.55±0.28 | 17.86 | 0.74-2.1 | **0.49** | **< 0.0001** |
|  | **2013** | 1.46±0.23 | 15.88 | 0.92-1.9 | **0.28** | **0.029** |
|  | **2014** | 1.37±0.24 | 17.66 | 0.86-2.38 | **0.35** | **0.0054** |
|  | **Mean** | **1.46±0.20** | 13.82 | 0.78-1.82 | **0.55** | **< 0.0001** |
| **FG** | **2011** | 114.49±38.69 | 33.80 | 63.50-284.2 | 0.14 | 0.430 |
|  | **2012** | 98.10±34.95 | 35.63 | 59.22-282.22 | 0.04 | 0.776 |
|  | **2013** | 141.56±33.66 | 23.78 | 70.07-220.71 | **0.32** | **0.014** |
|  | **2014** | 218.85±71.19 | 32.53 | 75.88-417.5 | **0.44** | **0.0005** |
|  | **Mean** | **143.25±30.83** | 21.52 | 67.75-246.14 | **0.38** | **< 0.0001** |
| **TG** | **2011** | 136.92±37.02 | 27.04 | 83.28-270.2 | 0.14 | 0.431 |
|  | **2012** | 115.26±34.96 | 30.33 | 82.30-288.95 | -0.02 | 0.870 |
|  | **2013** | 201.91±53.49 | 26.49 | 81.79-329.80 | **0.33** | **0.011** |
|  | **2014** | 253.05±87.34 | 34.51 | 82.57-517.7 | **0.37** | **0.0041** |
|  | **Mean** | 176.78±38.00 | 21.50 | 83.29-275.73 | **0.53** | **0.002** |
| **TGW** | **2011** | 24.05±3.27 | 13.59 | 14.77-32.0 | 0.17 | 0.239 |
|  | **2012** | 24.20±3.50 | 14.45 | 16.64-34.90 | **0.30** | **0.021** |
|  | **2013** | 22.85±3.01 | 13.18 | 15.24-29.02 | **-0.27** | **0.035** |
|  | **2014** | 26.25±4.61 | 17.57 | 17.08-38.58 | 0.02 | 0.8765 |
|  | **Mean** | 24.34±2.49 | 10.24 | 17.50-29.17 | **0.36** | **0.005** |
| **SLBR** | **2014** | 3.93±0.41 | 10.50 | 2.4-4.51 | 0.15 | 0.2481 |
|  | **Mean** | **3.57±0.41** | 11.55 | 2.44-4.52 | 0.17 | 0.566 |
| **YLD** | **2011** | 5.54±1.60 | 28.92 | **1.22-9.12** | 1.00 | **0** |
|  | **2012** | 5.63±1.61 | 28.56 | **1.73-9.89** | 1.00 | **0** |
|  | **2013** | 4.55±1.35 | 29.67 | 1.19-6.59 | 1.00 | **0** |
|  | **2014** | 5.90±1.68 | 28.41 | **1.95-9.6** | 1.00 | **0** |
|  | **Mean** | 5.41±1.48 | 27.33 | **1.82-8.80** | **1.00** | **0** |

Note: Significant level of alpha at 0.05% and $p < 0.0001$ ; Values in bold are different from 0 with a significance level alpha=0.05
DFF-Days to 50% flowering, PH-Plant height in cm, TL-Tiller number, PL-Panicle length in cm, FLL-Flag leaf length in cm, FLW-flag leaf width in cm, FG-No of fertile grains, TG-Total no of spikelets per panicle, TGW-1000-grain weight in gm, SLBR-Seed length-breadth ratio (SLBR), YLD-Grain yield t/ha.

Grain yield varied from 1.19 t/ha (Curinga, 2013) to 9.89 t/ha (N-129, 2014) with mean yield from 1.82 t/ha to 8.8 t/ha. Similarly, phenotypic variance for DFF varied from 72.5 days (Nipponbare, 2011) to 124.65 days (WC-8, 2014); 58.24 cm (Peta, 2012) to 186.93cm (WC-8, 2014) for PH; 5.0 (N-43, 2013) to 12.6 (N-49, 2013) for TL; 17.19 cm (Peta, 2012) to 34.8 cm (N-129, 2012) for PL; 17.72 cm (Peta, 2012) to 51.84 cm (WC-8, 2012) for FLL;, 0.65 cm (Peta, 2011) to 2.38 cm (N-224, 2014) for FLW; 59.22 (Peta, 2012) to 417.29 (N-373, 2014) for FG; 81.79 (Peta, 2013) to 517.7 (N-369, 2014) for TG; 14.77 gm (Sambamasuri, 2011) to 38.58 gm (N-65, 2014) for TGW and 2.4 (Peta) to 4.51 (N-318) for SLBR.

Some genotypes produced appreciably higher grain yield in respective seasons. Among them, N-129 produced highest grain yield in all four-season seasons (9.12 t/ha in 2011; 9.89 t/ha in 2012; 6.59 t/ha in 2013 (productivity affected due to Cyclonic Storm Phailin) and 9.60 t/ha in 2014). It was followed by R-261 (8.01 t/ha in 2011), N-370 (8.78 t/ha in 2012), N-8 (6.34 t/ha in 2013) and N-8 (8.86 t/ha in 2014), respectively. The mean of grain yield in four seasons varied from 1.82 to 8.8 t/ha. Twenty NPTs performed very well consistently in all the four seasons (S2 Table). The average grain yield of four seasons of these 20 genotypes varied from 5.45 to 8.8 t/ha, while popular varieties such as IR64 and MTU1010 could produce the maximum yield of 4.80 to 4.99 t/ha (Fig 1A, 1B and 1C, S2 Table).

**Fig. 1A, 1B, 1C.**
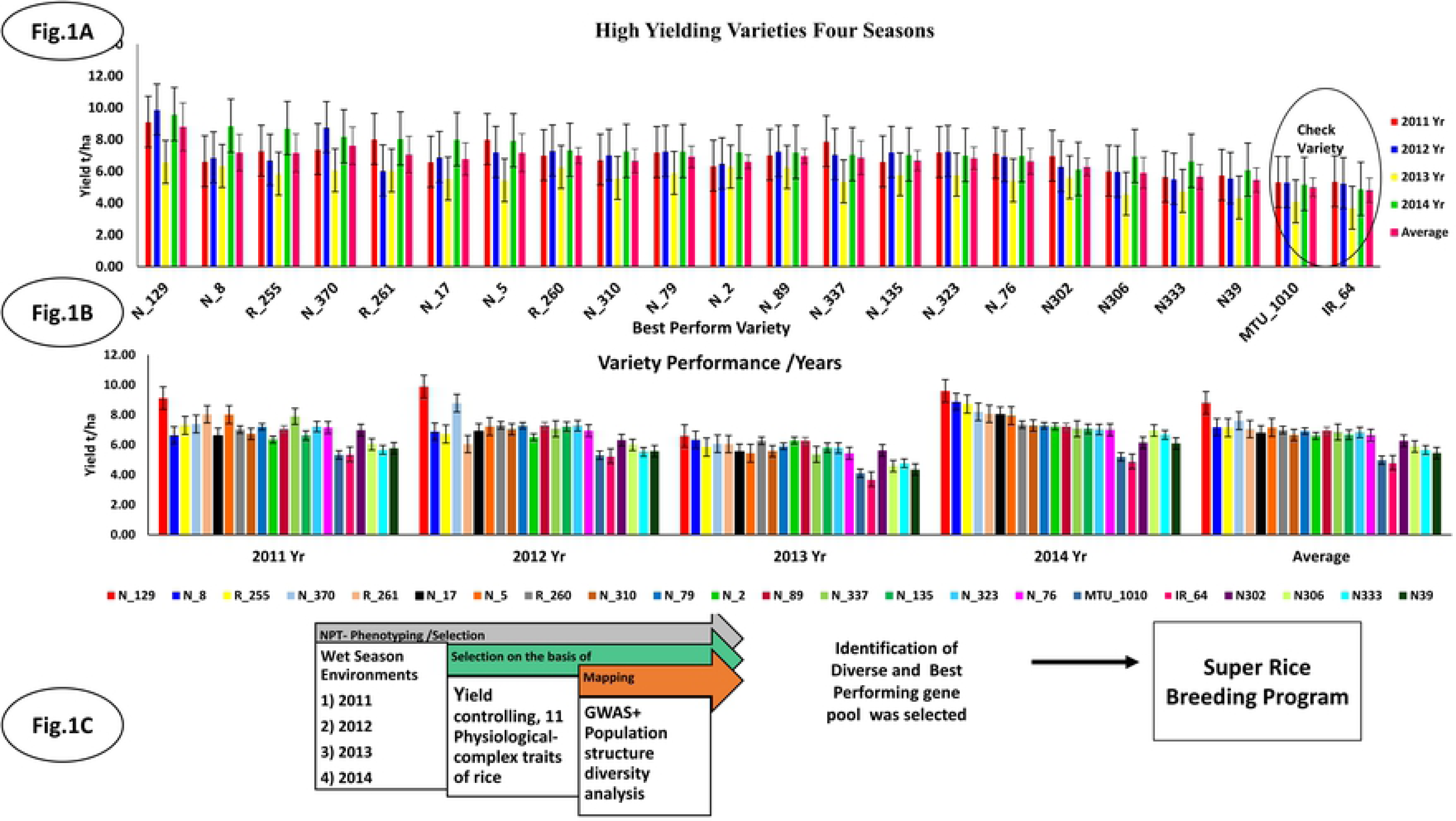
Performance of New Plant Types (NPTs) with reference to check varieties based on high grain yield in different seasons. The working hypothesis fulfil the objective of the study. Note: The circle of fig. (1-a) indicates the performance of standard check varieties i.e. IR-64 and MTU1010

The CV%, correlation and linear regression analysis of all traits were calculated at 5% level of significance (Table 1, S3 Table). Eight traits viz. DFF, PH, PL, TL, FLW, FG, TG and TGW were found to be positively correlated with mean of four-season grain yield (S3 Table). Similarly, linear regression showed a positive association of six yield contributing traits (PL, DFF, TL, FG, TG, FLL) with grain yield while four traits (PH, FLW, TGW, and SLBR) showed a negative association with grain yield (S4 Table, Fig 2A). The standardized coefficient plots showed that PH, SLBR and TG, were negatively associated grain yield, whereas DFF, TL, PL, FLL, FLW and FG were positively associated with grain yield (S4 Table, Fig 2A). The standardized plot also showed that PH, SLBR, TG and TGW were not directly involved in controlling grain yield. Their significance level may be influenced by environmental factors. The standardized coefficient plot was shown with positive bars for genotypes, which showed grain yield of more than 6.0 t/ha. Similarly, a negative bar of traits having less grain yield is not associated with traits (S4 Table, Fig 2A).

**Fig 2A.**
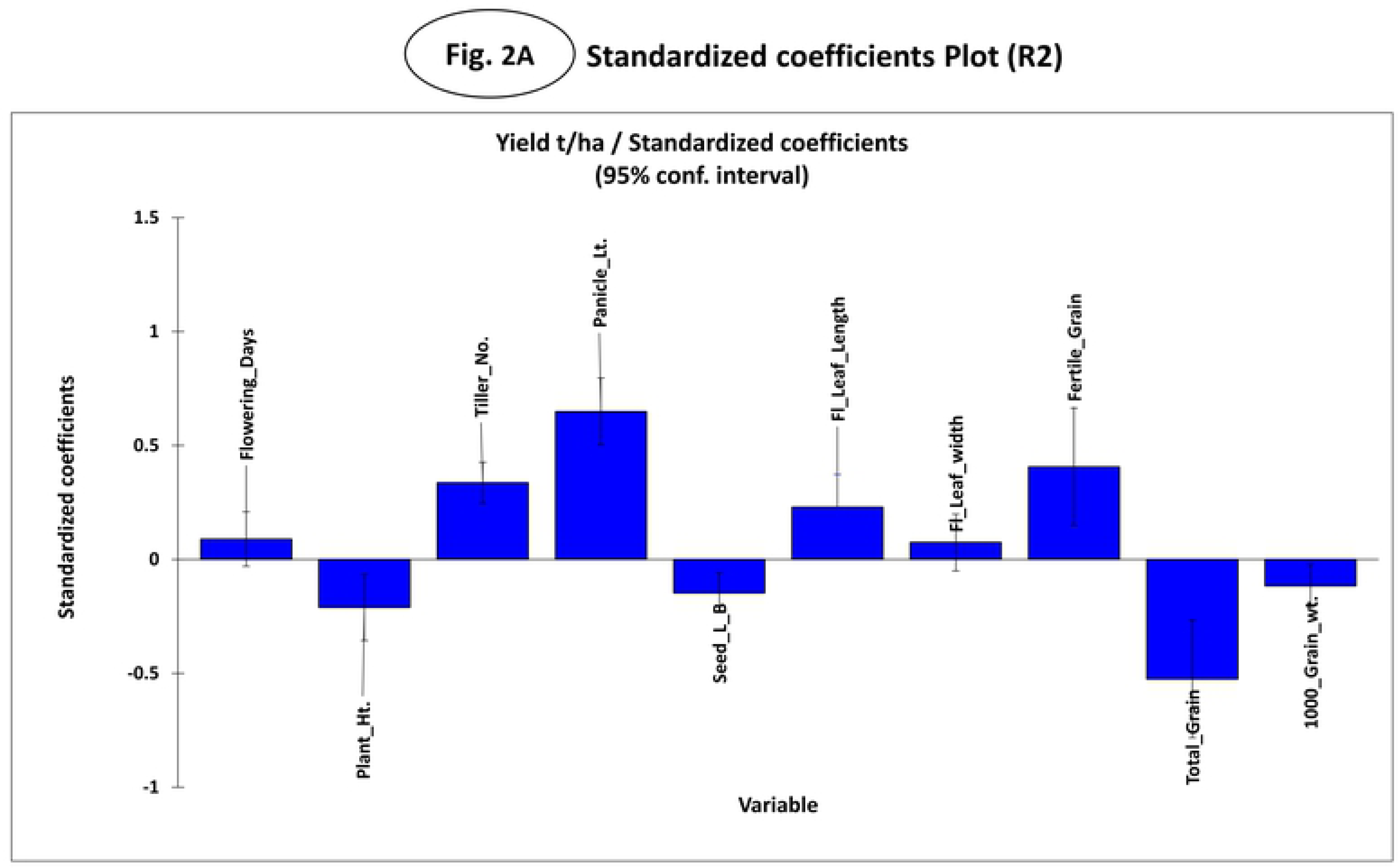
The standardized coefficient plot has shown the correlation between yield and yield-related traits and genotypes in a regression analysis. The positive bar indicates having a positive correlation with the traits and vice versa for the negative bar.

**Fig 2B.**
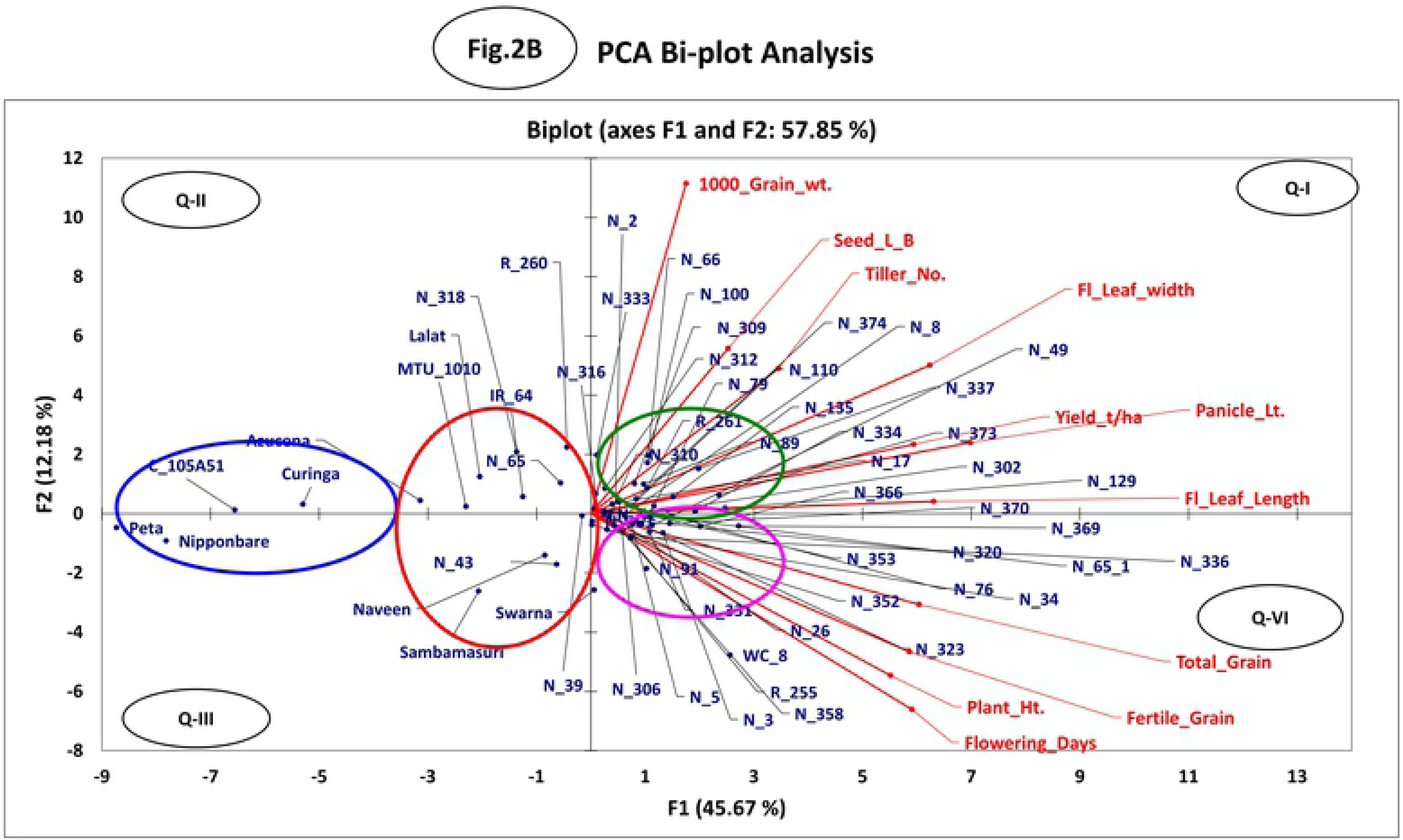
In PCA Biplot analysis, the genotypes were scattered along with their similarity and performance.

The PCA Biplot analysis was carried out for focusing on dominant phenotypic traits (Fig 2B). The first PC1 explained 45.67 %, while PC2 explained an additional 12.18 % of the phenotypic variance. The analysis indicated that the traits viz., TGW, SLBR, TL, FLL, FLW, PL and YLD were predominant for the genotypes situated in the green circle in quadrant I (S5 Table). Similarly, the genotypes belonging to the pink circle were having a preponderance of traits viz., TG, FG and PH. However, the trait DFF was dominant in quadrant IV (lower right side) (Fig 2B).

The circle represents genotypes are close to each other and have many similarities between them. The green and pink circles were having the genotypes that produced approximately 6-10 t/ha grain yield. The Red and Blue circles were representing genotypes having less grain yield, i.e. 3-4 t/ha and it was distinguished from right-side circles. The present study reported a broad range of grain yield in different years, viz., 1.22-9.12 t/ha in 2011; 1.73-9.89 t/ha in 2012; 1.19-6.59 t/ha in 2013; 1.95-9.6 t/ha in 2014, whereas the mean of four seasons was found to be 1.82-8.80 t/ha (Table 1, Fig 1). The current study identified sixteen NPT genotypes, which were performed better than standard check variety IR64 and MTU1010 (Fig 1). The three best NPT genotypes were identified having highest grain yield in N-129 i.e. 9.12 t/ha (WS 2011), 9.59 t/ha (WS 2012), 6.59 t/ha (WS 2013) and 9.6 t/ha (WS 2014), respectively. N-8 showed second highest grain yield in four seasons i.e. 6.63 t/ha (2011), 6.88 t/ha (2012), 6.34 t/ha (2013) and 8.86 t/ha (2014). Similarly, third-highest grain yield was reported for R-255 i.e., 7.30 t/ha (2011), 6.72 t/ha (2012), 5.85 t/ha (2013) and 8.72 t/ha (2014) in four consecutive years. However, these genotypes could be used as a donor for yield-related specific traits.

Heat map helps to understand diversity and dominance pattern between genotypes and traits. Heatmap showed that dominant traits were grouped into two significant clusters of genotypes basing on certain similarities. In traits, the first cluster included FG, TG, YLD, TL, FLW, and SLBR, whereas the 2^nd^ cluster comprised TGW, PH, FLL, DFF and PL, which were found very important for specific genotypes (Fig 3). In map red colour traits, i.e., FG, TG, PH and DFF in clusters shows their dominance for respective genotypes. The trait, total grain number (TG) is dominant in most of the NPT genotypes which belongs in 2^nd^ cluster (Fig 3).

**Fig 3.**
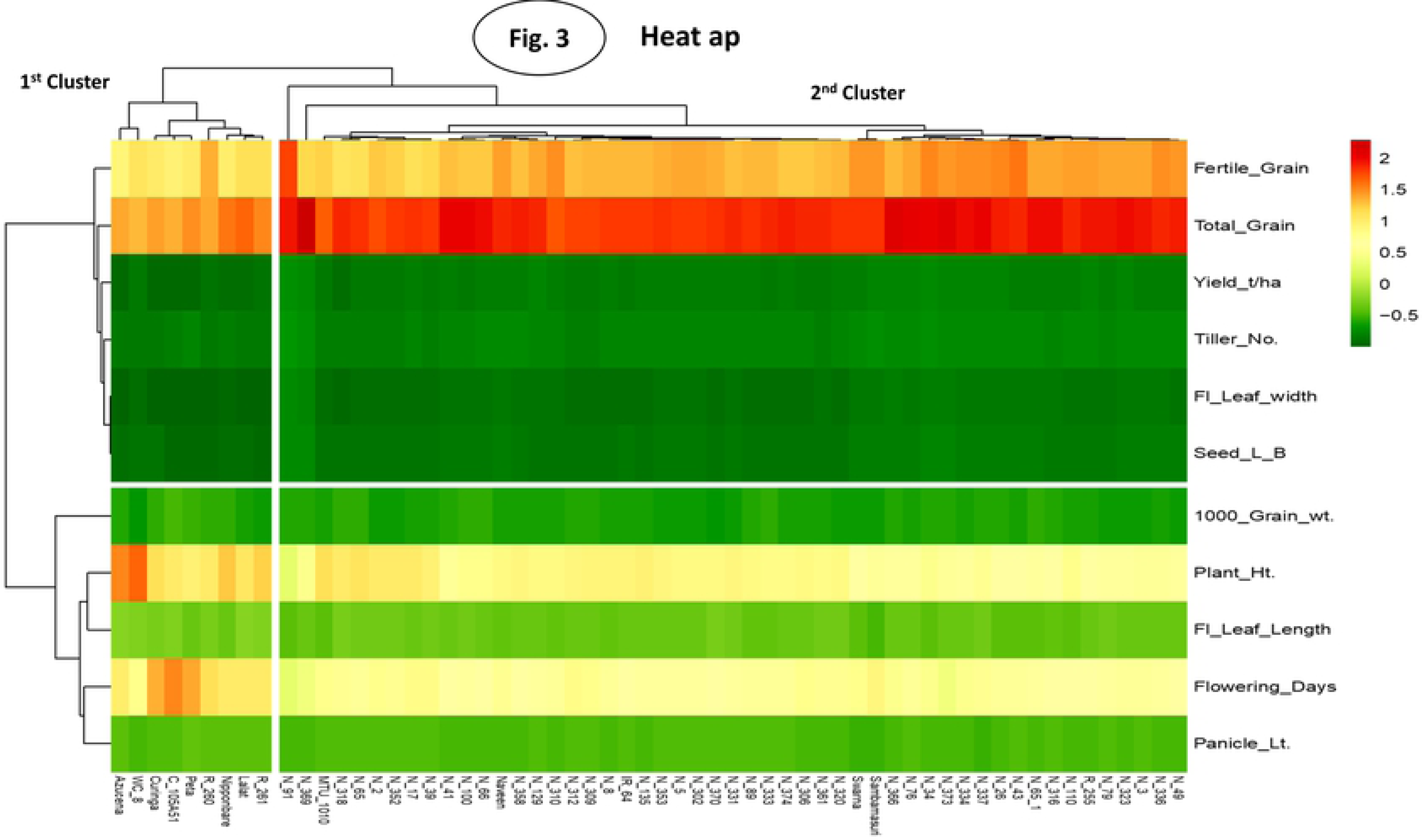
Heat Map showing that both genotypes and traits are grouped into rows and columns by using correlation distance and average matrix.

### Allelic Diversity

Sixty-six (77.65%) out of 85 SSRs were found to be polymorphic. A total of 154 alleles were amplified by 66 polymorphic microsatellite markers with an average of 2.33 alleles per locus. Five markers viz., RM154, RM5709, RM204, RM70 and RM1132 amplified the highest number of alleles (i.e., 4). Two unique alleles (amplified by RM6266 and RM489) and eight rare alleles (5.19%) were identified (Table 2). The marker RM5709 amplified two rare alleles while markers RM168, RM6266, RM489, RM3276, RM528, and RM70 amplified one rare allele each (S6 Table). The major allele frequency (MAF) varied from 0.33 (RM1132) to 0.98 (RM6266) with an average frequency of 0.71. The genetic diversity varied from 0.033 (RM6266) to 0.732 (RM1132) with an average of 0.39 per locus. The number of effective alleles (Ne) varied from 1.034 (RM489) to 3.733 (RM1132) with an average of 1.78 per locus. The homozygosity (Ho) ranged from 0.262 (RM1132) to 0.967 (RM6266) with an average of 0.60, while genetic heterozygosity (He) varied from 0.033 to 0.732. Shannon’s information/diversity index (*I*) ranged from 0.085 (RM6266) to 1.347 (RM1132) with an average of 0.39 and 0.63 per locus for both the parameters, respectively. The polymorphism information content (PIC) indicates the allelic diversity and frequency of a marker locus with respective genotypes. It varied from 0.516 (RM6266 and RM489) to 0.92 (RM204) with an average of 0.70 (S6 Table). Positive correlations were observed between the total number of alleles (TA), low-frequency alleles (LFA), high-frequency alleles (LFA) and PIC (S6 Table, S7 Table).

**Table 2.** Unique and rare alleles amplified by microsatellite loci in different rice genotypes.

| Chromosome No. | Primers | bp | Genotypes |
| --- | --- | --- | --- |
| **a)Unique allele** |  |  |  |
| 3 | RM 6266 | 160 | N-361 |
| 3 | RM 489 | 300 | N-34 |
| **b)Rare alleles** |  |  |  |
| 3 | RM168 | 80 | Lalat, Samba Mahsuri |
| 3 | RM 6266 | 160 | N-361 |
| 3 | RM 489 | 300 | N-34 |
| 4 | RM 3276 | 190 | N-334, N-110, N-318 |
| 4 | RM 5709 | 200 | C105A-51, AC41009, Nipponbare, N-48 |
| 4 | RM 5709 | 220 | N-2, N-333, N-320 |
| 6 | RM 528 | 260 | MTU1010, Samba Mahsuri, R-260 |
| 7 | RM 70 | 200 | R-266, N-306 |

### Genetic Diversity

The Nie’s pairwise genetics analysis by Neighbour-joining tree grouped all the 60 genotypes into three clusters/populations (POP1, POP2 and admixture) (Fig. 4A). The first cluster includes four sub-clusters containing *Indica*, *Trop. Japonica, Tem. Japonica* and one NPT genotypes were identified. However, second and third clusters included 47 NPT genotypes, some of which found as an admixture in the second cluster. The NPT genotypes have been derived from *indica, tropical* and *temperate japonica*. Hence, two types of populations were observed along with admixture cluster. Both the populations in the trees were observed to be distinctly different (Fig 4A, Table 3, S6 Table).

**Fig 4A, 4B.**
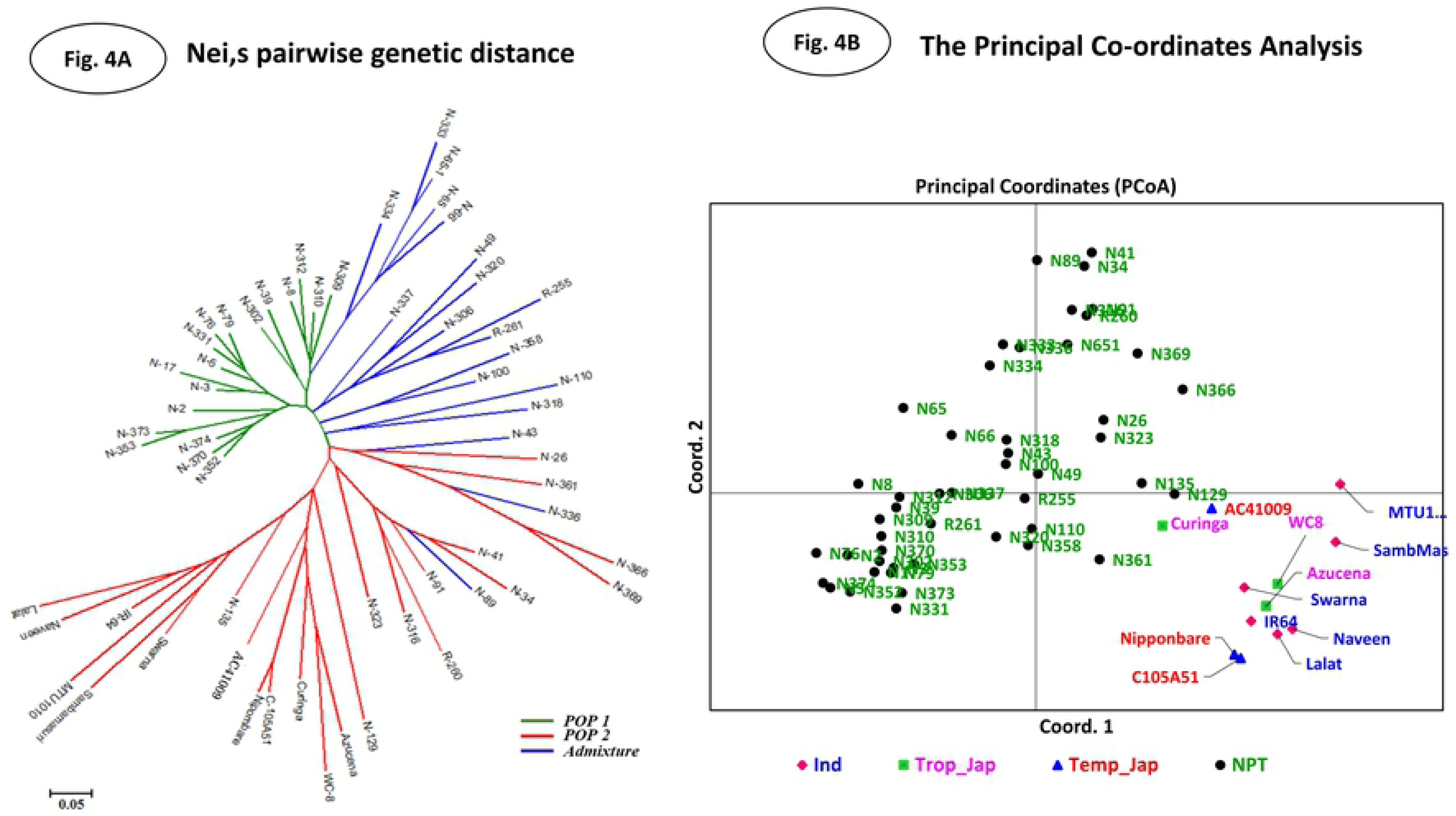
The pairwise genetic distance (Nei, 1973) was calculated by POPGENE v 1.32, and it shows genotypes distributed according to their archetypes. The Principal Coordinate analysis (PCoA) of 60 genotypes based on 66 SSR markers. The graph shows the position, and the distribution pattern of each genotype in population space spanned by coordinate 1^st^ versus coordinate 2nd.

**Table 3.**
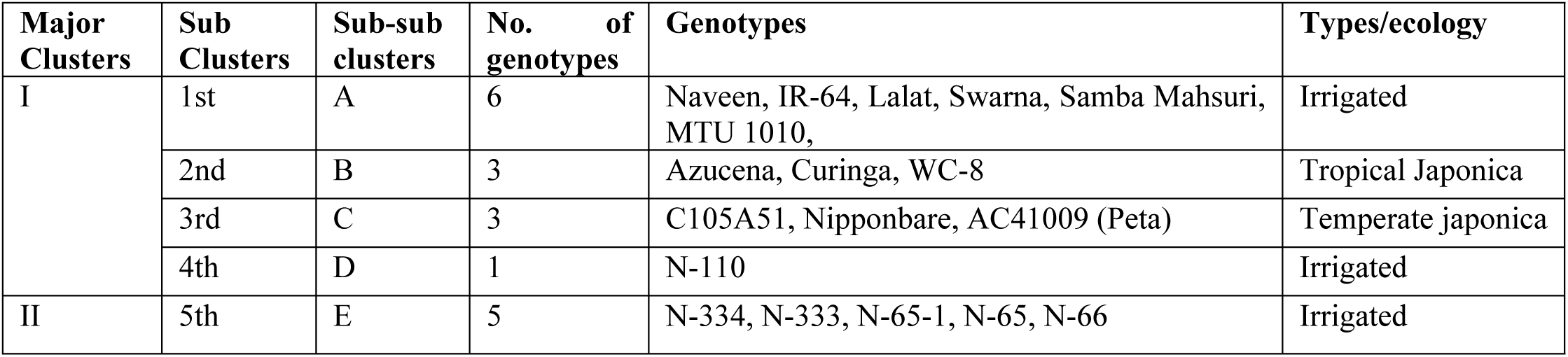
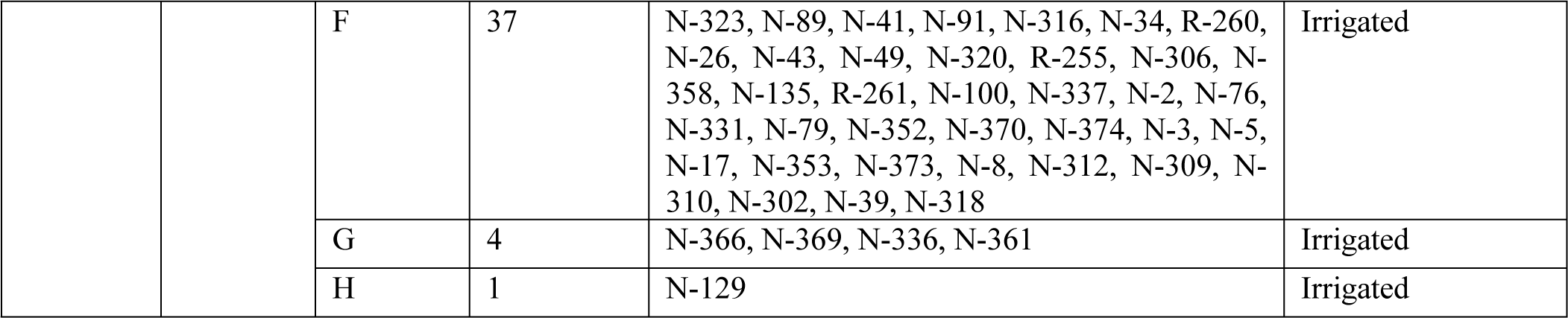
Grouping of 60 genotypes based on the UPGMA analysis.

The Principal Coordinate Analysis (PCoA) differentiated all the 60 genotypes and separated NPTs from *indica* and *japonica* genotypes (Fig 4B). *Japonica* and *indica* genotypes were grouped separately and slightly different from each other. However, many NPT genotypes were grouped into the separate cluster as per neighbour-joining cluster and structure analysis. The PCoA percentage of molecular variance explained by three axes was found to be 12.43%, 7.93%, and 7.45%. In PCoA, the 4^th^ quadrant group showed that some NPTs having an admixture of *indica* and *japonica* populations were intermixing (Fig 4B).

The UPGMA cluster analysis grouped all the 60 genotypes into two major clusters at 54% genetic similarity. The first major cluster (I) was sub-grouped into four sub-clusters, i.e., A, B, C, and D with similarity index varied from 0.54 to 0.92. These sub-clusters contained all the six *indica*, three tropical *japonica*, and three temperate *japonica* and one NPT, respectively (Fig 5, Table 3). Second major cluster (II) contained 47 NPTs with similarity index varied from 0.56 to 0.91. Further, it was sub-grouped into four sub-clusters i.e., E, F, G, and H containing 5, 37, 4 and 1 genotypes, respectively. The sub-cluster D and H contained only one genotype each, i.e., N-110 and N-129, respectively (Fig 5, Table 3).

**Fig 5.**
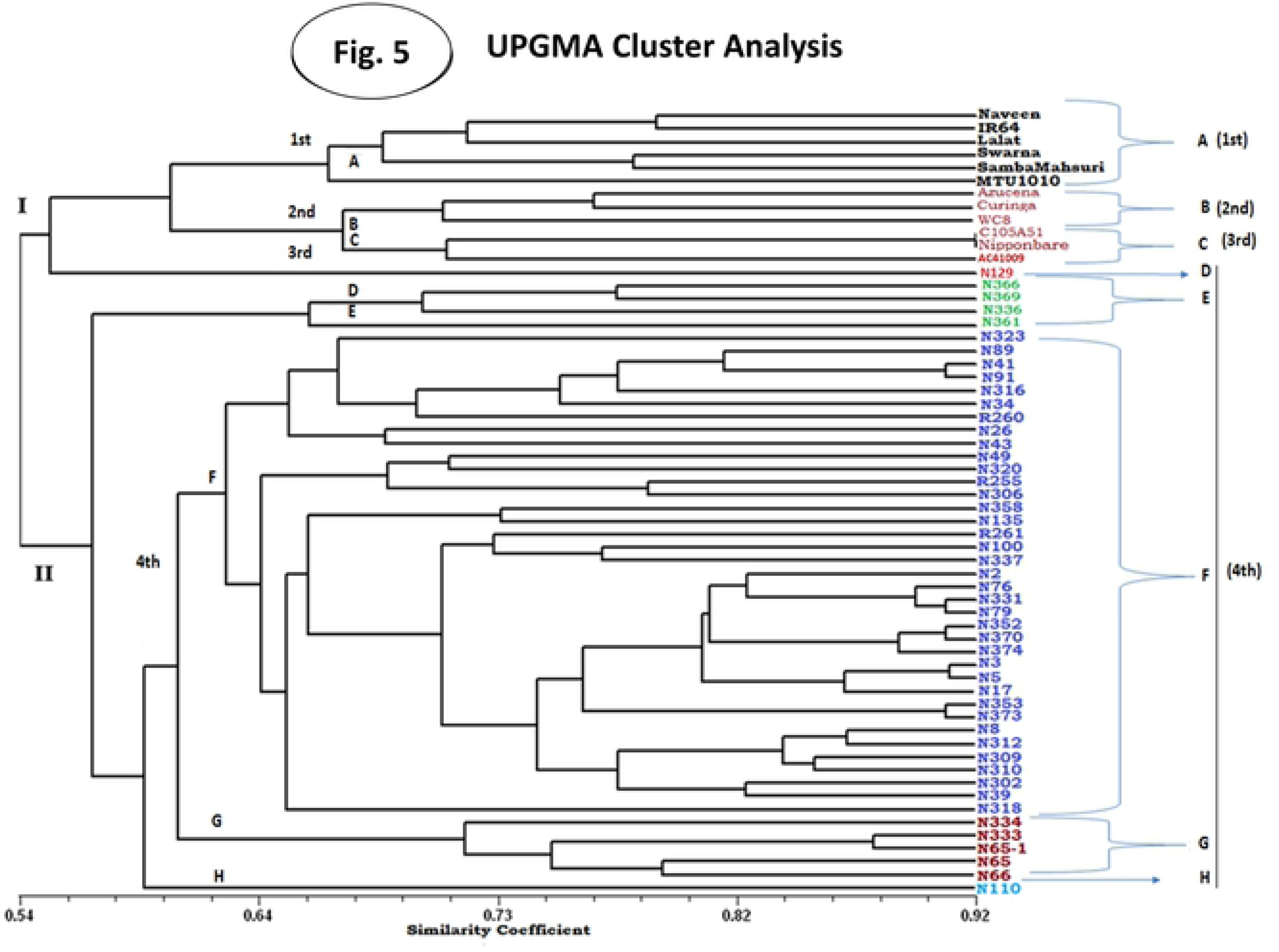
The genotypes clustering based on UPGMA analysis. The 60 genotypes were assigned into four groups (A-1st, B-2nd, C-3rd, and D, E, F, G, H-4th) and similarity varied between 0.54 to 0.92%.

### Analysis of Molecular Variance (AMOVA)

The two populations obtained through STRUCTURE analysis were used to know the genetic variation between and within clusters using AMOVA. The analysis accounted for 8% within individuals (genotypes), 67% among individuals within a population and 25% among populations (Fig 6, Table 4). The F-statistics on all three groups found to be highly significant (P<0.001). The overall Fst (Fst=0.240) had significant (P<0.001) genetic variation among the three populations (Fig 6, Table 4).

**Fig 6.**
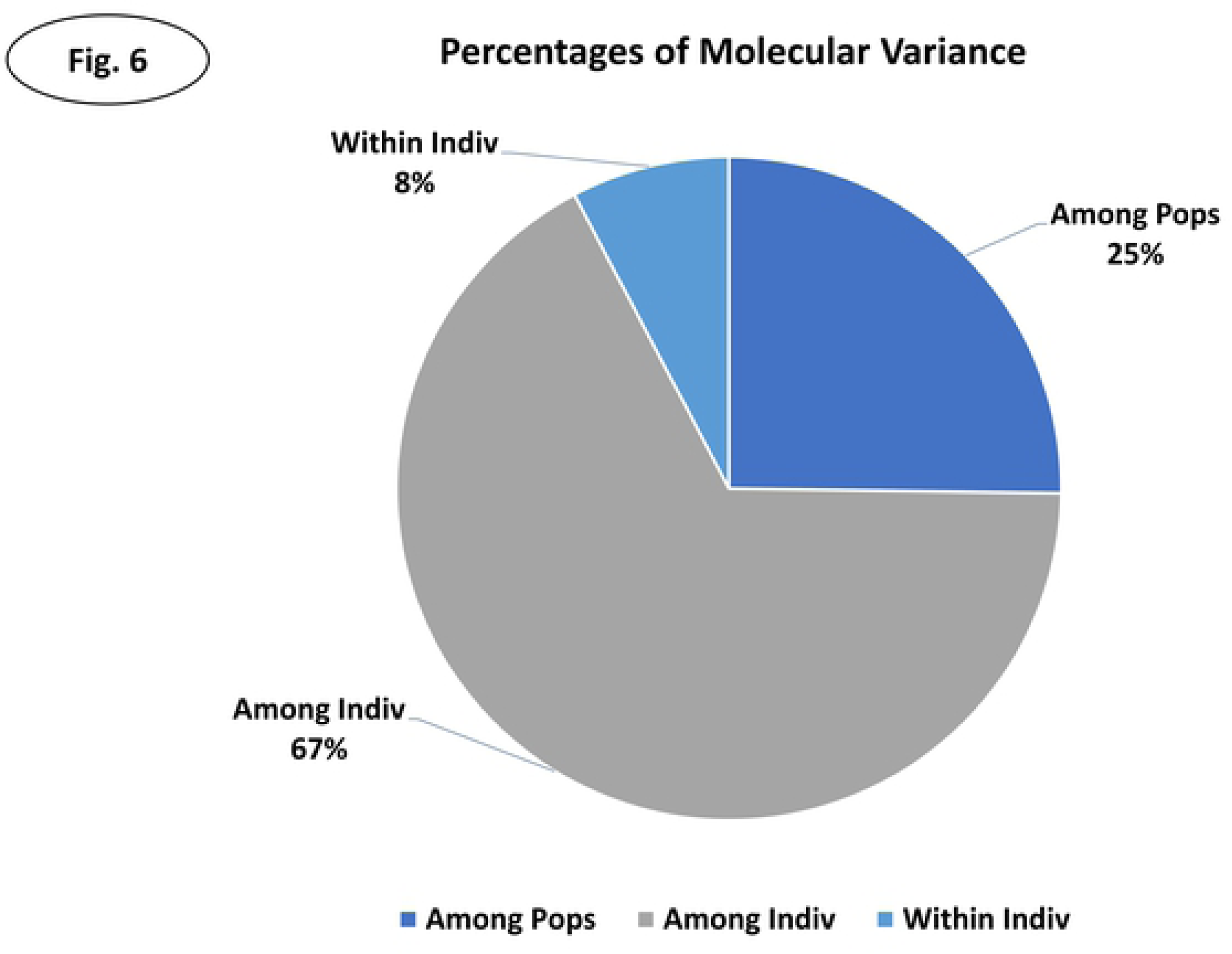
AMOVA-showing molecular distribution pattern within and among populations.

**Table 4.** Analysis of molecular variance (AMOVA) among and between populations

| Source of variation | df | SS | MS | Estimated Variation | % variation of | F statistics | Value | P value |
| --- | --- | --- | --- | --- | --- | --- | --- | --- |
| Among Populations | 2 | 198.9 | 99.5 | 3.8 | 24% | $F_{st}$ | 0.240 | <0.001 |
| Among Individuals within populations | 57 | 1288.8 | 22.6 | 10.7 | 68% | $F_{is}$ | 0.898 | <0.001 |
| Within Individuals (genotypes) | 59 | 73.0 | 1.3 | 1.3 | 8% | $F_{it}$ | 0.922 | <0.001 |
| Total | 118 | 1560.7 |  | 15.7 | 100% |  |  |  |

**MS**-Mean sum of square
**$F_{st}$** - F-statistics; subpopulations within the total population
**$F_{is}$** - F-statistics; individuals within subpopulations

### Population Structure

The true value of K was identified according to the maximum value of LnP (D) (Pritchard et al., 2000). Structure harvester of Evano table (http://taylor0.biology.ucla.edu) analysis showed that at K=2, the ΔK=179.57, where value was the highest in both independent burns [40]. The ΔK values were decreased from K= 2 to 10 in general but had a moderate value of 56.29 at K=4. At K=4, all the 60 genotypes were divided into four subpopulations, POP1, POP2, POP3, and POP4, which contained 6 *indica*, 3 temperate *japonica*, 3 tropical *japonica*, and 48 NPT genotypes, respectively (Fig 7). The populations POP2, POP3, and POP4 showed admixture types. Both Pritchard’s and Evonne’s methods confirmed the K-value as 2. Furthermore, analysis of POP gene software showed that 60 genotypes was grouped into two major populations, followed by one admixture group (Fig 7, S1 Fig). The STRUCTURE analysis genotypes were grouped into two types of populations at K=2, while at K=3 and K=4, 60 genotypes were grouped into three and four types of populations, respectively.

**Fig 7.**
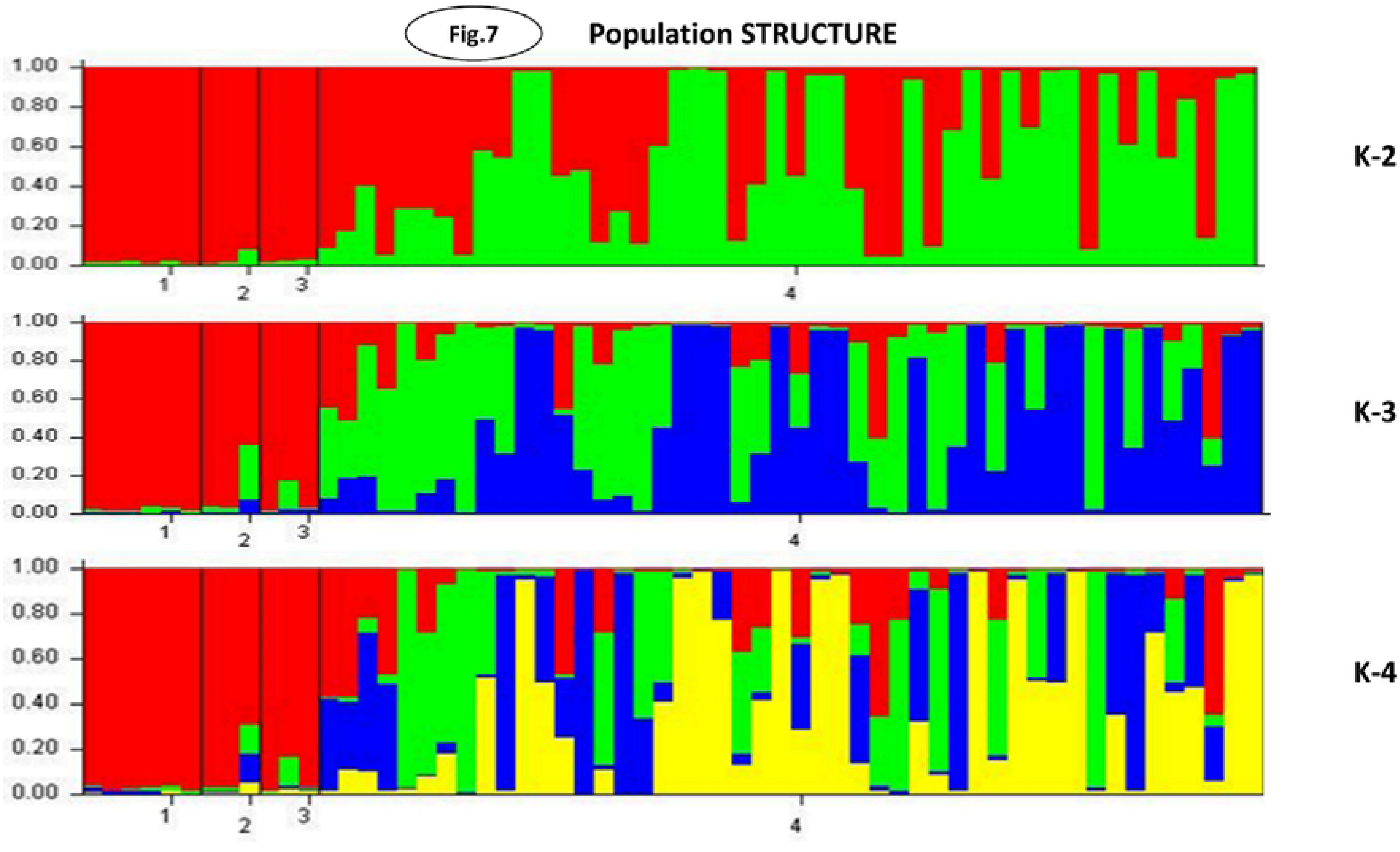
The true value of K was determined by STRUCTURE harvester in K-2 Plot of change in the likelihood of the data, L(K), at values of K from 1 to 10 K=2.

### QTL Association

#### Marker-trait in general

The GLM and MLM models were used for association analysis (p-value <0.005 and FDR at 5% level. A total of 31 SSR markers were found to be associated with grain yield and other 10 related traits based on individual seasons and their mean data using GLM and MLM models (Table 5, Fig. 8). A total 30 and 10 SSRs identified by GLM and MLM models were associated with 70 and 16 QTLs, respectively. Fifteen SSRs (RM6100, RM1132, RM222, RM297, RM154, RM168, RM551, RM5709, RM5575, RM20285, RM5711, RM234, RM26499, RM19 and RM204) were identified to be associated with grain yield *per se* (YLD). Similarly, association analysis led to the identification five SSRs with DFF, three each with PH, FG and TG, four each with TL, FLL and SLBR, seven SSRs with TGW, 13 SSRs with PL and 14 SSRs with FLW (Table 5, Fig 8).

**Fig 8:**
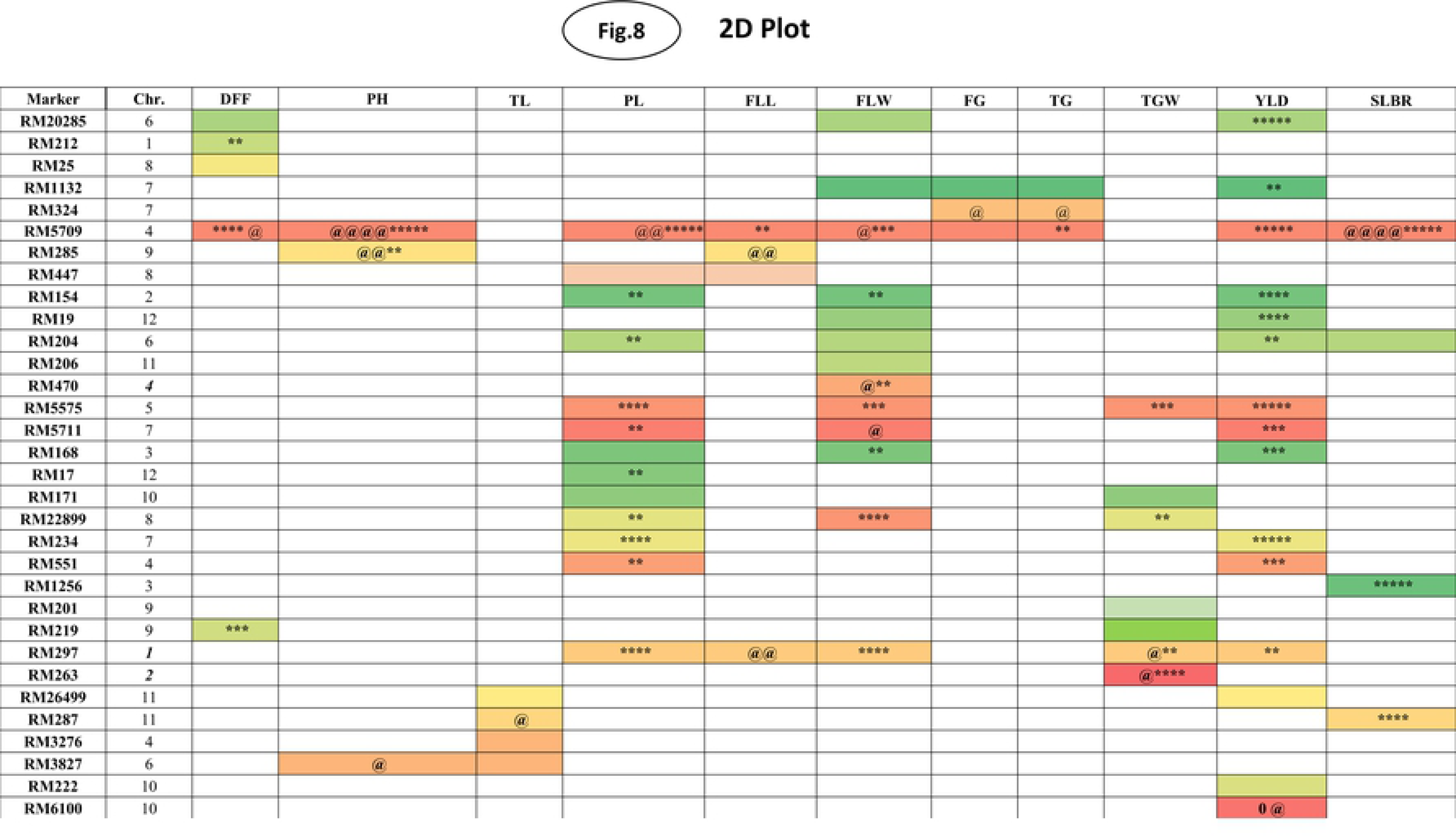
The 2D-Plot showing SSR marker association with respective phenotypic traits using GLM (p-0.005) and MLM (p-0.005) models. **MLM***=**@** -Markers associated with MLM in the specific -season /environment* *\*\*/ @@= Indicates SSR markers associated in two seasons/ environments* *\*\*\*/ @@@= Indicates SSR markers associated in three seasons/ environments* *\*\*\*\*/ @@@@= Indicates SSR markers associated in four seasons/ environments*

**Table 5.**
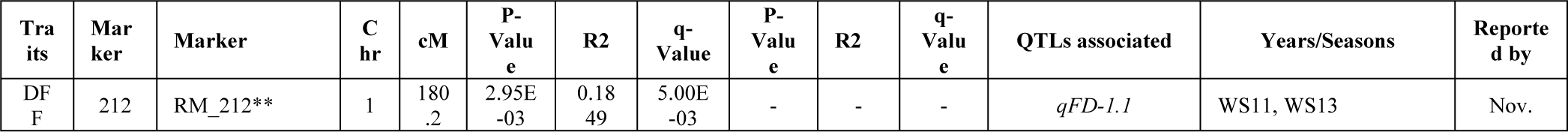
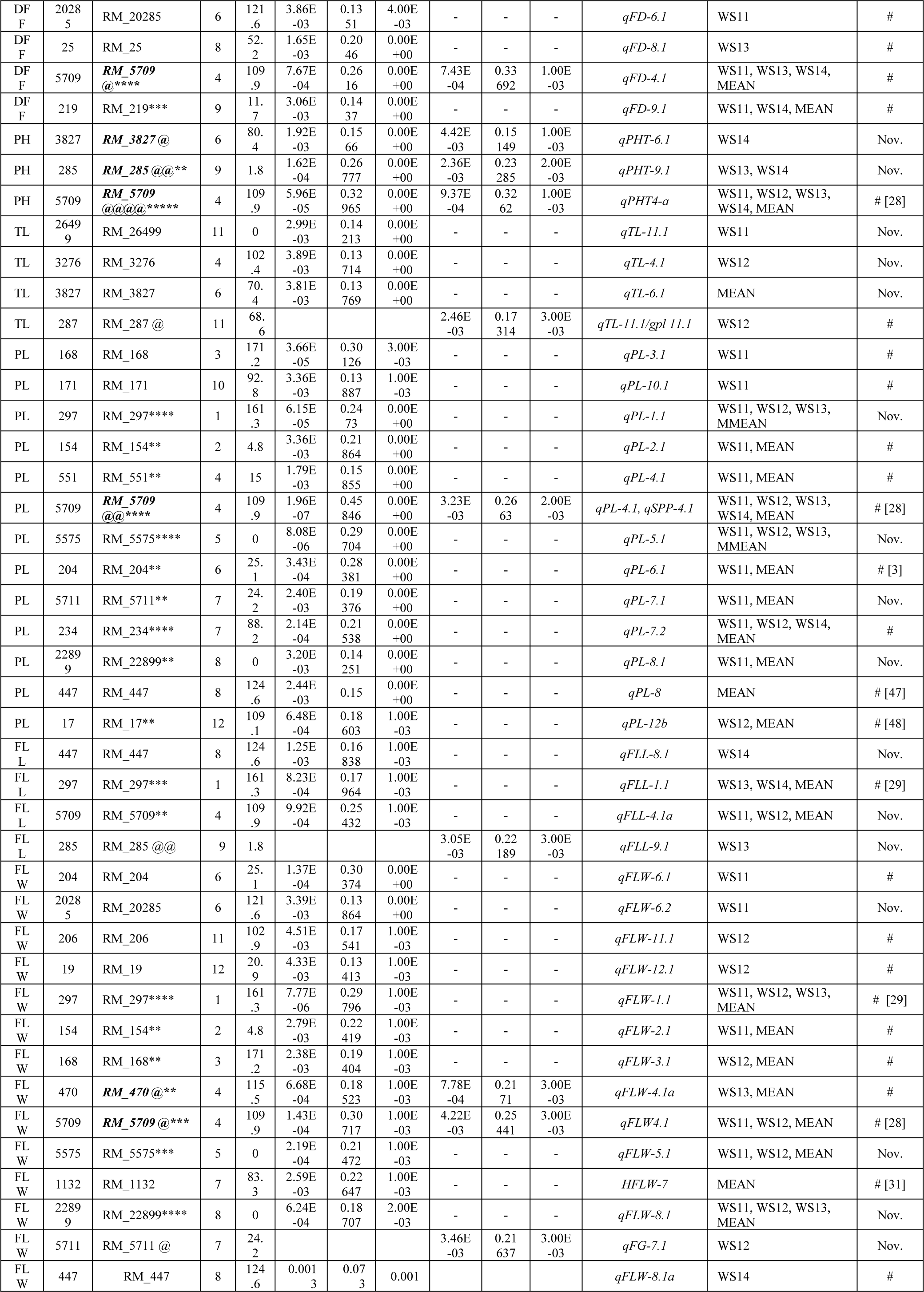
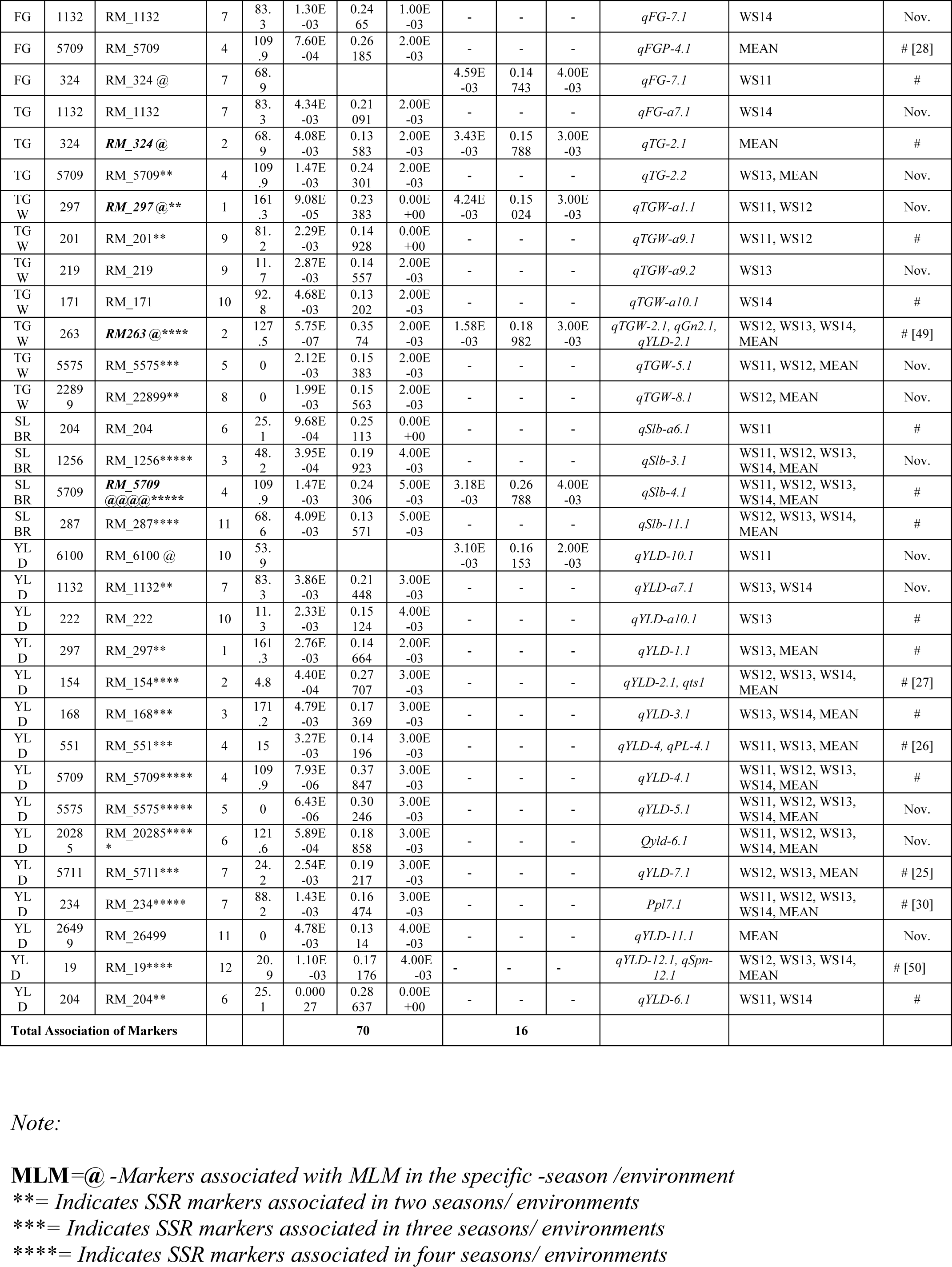
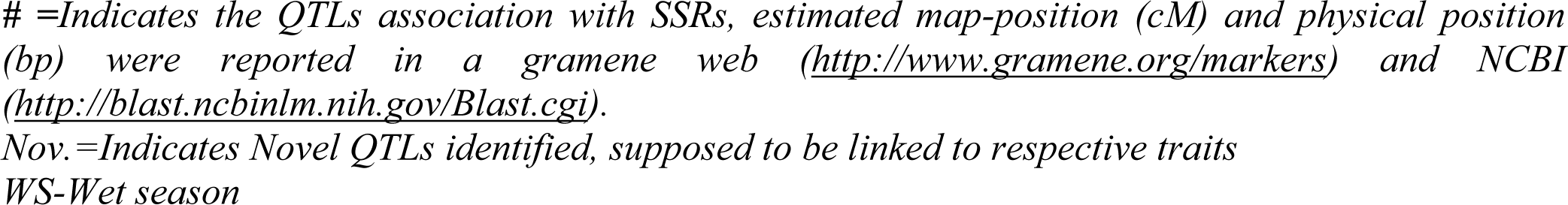
Identification of SSR markers associated with grain yield and yield-related traits by using GLM and MLM models.

The MLM analysis identified 10 SSRs that are significantly associated (p-value <0.005) with 16 QTLs, which were association with six traits based on the mean data of four seasons at 5% FDR (Table 5). Four SSRs were found to be associated with QTLs considering four seasons data separately as well as their mean data (viz., *qPHT4-a*; *qPL-4.1* (*qSPP-4.1*); *qFLW-4.1* and *qTGW-2.1* (*qTGW-2.1, qGn2.1, qYLD-2.1*). The SSR marker RM5709 was found to be associated with nine traits, i.e. DFF (*qFD-4.1*), PH (*qPHT4-a*), PL (*qPL-4.1*/ *qSPP-4.1*) FLL, FLW (*qFLW-a4.1* and *qFLW-4.1*), FG (*qFGP4.1*), TG (*qTG2.2*) SLBR (*qSlb-4.1*) and YLD (*qYLD4.1*) indicating its association with these major traits. Three markers RM285, RM3827 and RM5709 were found to be associated with QTLs for plant height, *qPHT-6.1*, *qPHT-9.1* and *qPHT4-a*, respectively (Table 5; Fig 8). Similar marker-QTL association was also recorded for complex traits like TGW (*qTGWa1.1* and *qTGW-2.1*, *qGn2.1, qYLD-2.1*), FLL (*qFLL-9.1*), TG (*qTG-2.1*) and FG (*qFG-7.1*) (Table 5). As two different models have different association with respective traits and therefore, reliability of marker-traits, association would be on the basis of the number of times it was showing association with respective traits in different seasons.

The phenotypic variance contributed by QTLs/SSRs found to be 13.51 % (RM20285) to 26.16 % (RM5709, *qFD4.1*) for DFF, 15.15 % (RM3827, *qPHT6.1*) to 32.96 % (RM5709, *qPHT4a*) for PH, 13.71 % (RM3276, *qTL4.1*) to 17.34 % (RM287, *qTL11.1/qgpl11.1*) for TL, 13.89 % (RM171) to 45.85 % (RM5709, *qPL4.1, qSPP4.1*) for PL, 16.84 % (RM447, *qFLL8.1*) to 25.43 % (RM5709) for FLL, 0.7 % (RM447, *qFLW-8.1a*) to 30.72 % (RM5709, *qFLW4.1*) for FLW, 14.74 % (RM324, *qFG7.1*) to 26.19 % (RM5709, *qFGP4.1*) for FG, 13.58 % (RM324, *qTG2.1*) to 24.30 % (RM5709, *qTG2.2*) for TG, 13.20 % (RM171, *qTGWa10.1*) to 35.74 % (RM263, *qTGW2.1, qYLD2.1, qGn2.1*) for TGW and 13.57 % (RM287, *qSlb11.1*) to 26.79 % (RM5709, *qSlb4.1*) for SLBR and 13.14 % (RM26499, *qYLD11.1*) to 37.85 % (RM5709, *qYLD4.1*) for yield (Table 5; Fig 8). More than 25% phenotypic variance was explained by QTLs bracketing RM5709 for each of nine traits, DFF, PH, PL, FLL, FLW, FG, TG, SLBR and YLD. This indicated that RM5709 would be useful for transfer above nine traits into popular rice varieties.

Twenty SSR markers were found to be significantly associated with more than one trait (Table 6). RM5709 was found to be associated with nine traits while RM275 was found to be associated with five traits. Similarly, RM5575, RM204 and RM1133 were found to be associated with four traits each while RM154, RM168, RM20285, RM5711, RM447, RM22899 were found to be associated with three traits each. Nine SSR markers were found to be associated with two traits (Table 6).

**Table 6.** Identification of SSR markers associated with more than one trait

| Marker | Chromosome | Position | Co-localization Traits | Gene ID | Locus ID | Function | Reference |
| --- | --- | --- | --- | --- | --- | --- | --- |
| RM_297 | 1 | 32,099,566-32,099,760 | PL, FLL, FLW, TGW, YLD |  |  |  | [51] |
| RM_154 | 2 | 1,083,820-1,084,090 | PL, FLW, YLD | Os02g0120800 | LOC_Os02g02840.1 | Similar to Small GTP-binding protein, EST Rice root cDNA, mRNA | [51–53] |
| RM_168 | 3 | 28,091,534-28,091,727 | PL, FLW, YLD |  |  |  | [51,54] |
| RM_5709 | 4 | 31,875,481-31,875,663 | DFF, PH, PL, FLL, FLW, FG, TG, YLD, SLBR | Os04g0619500 | LOC_Os04g52850.1 | Ovarian tumour, otubain domain containing protein. | [51–53] |
| RM_551 | 4 | 177,080-177,271 | PL, YLD |  |  |  | [51,54] |
| RM_5575 | 5 | 23,345,264-23,345,401 | PL, FLW, TGW, YLD | Os05g0474600 | LOC_Os05g39680.1, LOC_Os05g39690.1 | Similar to Aldose reductase-related protein | [51–53] |
| RM_20285 | 6 | 22,420,889-22,421,088 | DFF, FLW, YLD | Os06g0561800 | LOC_Os06g36650.1 | ABC transporter, transmembrane domain containing protein | [51–53] |
| RM_204 | 6 | 3,168,314-3,168,547 | PL, FLW, YLD, SLBR | Os06g0162800 | LOC_Os06g06750.1 | MADS-box transcription factor, Control of spikelet morphogenesis, Regulation of floral meristem | [51–53] |
| RM_3827 | 6 | 22,297,146-22,297,320 | PH, TL | Os06g0560300 | LOC_Os06g36480.1 | Similar to NAC-type transcription factor. | [51–53] |
| RM_1132 | 7 | 23,984,489-23,984,581 | FLW, FG, TG, YLD |  |  |  | [51] |
| RM_324 | 7 | 11,389,704-11,389,942 | FG, TG |  |  |  | [51] |
| RM_5711 | 7 | 3,141,181-3,141,329 | PL, FLW, YLD | Os07g0157700 | LOC_Os07g06390.2, LOC_Os07g06390. | Conserved hypothetical protein. | [51–53] |
|  |  |  |  |  | 3,<br>LOC_Os07g06390.<br>4 |  |  |
| RM_234 | 7 | 25,472,688-25,472,820 | PL, YLD |  |  |  | [51] |
| RM_447 | 8 | 26,546,992-26,547,102 | PL, FLL,<br>FLW | Os08g053120<br>0 | LOC_Os08g41900.<br>1 | WD40 repeat domain<br>containing protein | [51–53] |
| RM_22899 | 8 | 14,762,955-14,763,118 | PL, FLW,<br>TGW | Os08g033190<br>0 | LOC_Os08g24310.<br>1 | Similar to Leucine Rich<br>Repeat family protein,<br>expressed | [51–54] |
| RM_285 | 9 | 1.8-1.8 cM | PH, FLL |  |  |  | [51] |
| RM_171 | 1<br>0 | 19,048,795-19,049,123 | PL, TGW | Os10g048990<br>0 | LOC_Os10g34820.<br>1,<br>LOC_Os10g34820.<br>2 | Similar to CDT1a<br>protein | [51–53] |
| RM_26499 | 1<br>1 | 11,001,140-11,001,342 | TL, YLD |  |  |  | [51] IRGSP-1 |
| RM_287 | 1<br>1 | 16,767,319-16,767,617 | TL, SLBR |  |  |  | [51] IRGSP-1 |
| RM_19 | 1<br>2 | 20.9-20.9<br>cM | FLW, YLD |  |  |  | [51]/IRGSP-1 (2005) |

Nineteen SSR markers were found to be associated with different traits in more than one season. Four SSRs i.e., RM5709, RM5575, RM20285 and RM234 were found to be associated with PL, PH, SLBR and YLD and common for 4 seasons. Among them, RM5709 is co-localized with PH, PL, SLBR and YLD in four seasons. Ten SSRs were co-localized in three seasons with seven traits i.e., RM5709 (DFF, PH), RM297 (PL), RM5575 (PL), RM234 (PL), RM22899 (FLW), RM263 (TGW), RM1256 (SLBR), RM287 (SLBR), RM154 (YLD) and RM19 (YLD). Similarly, 12 SSRs were found to be associated with two seasons viz., RM212 (DFF), RM219 (DFF), RM285 (DFF), RM297 (FLL, TGW), RM5709 (FLW, FLL), RM201 (TGW), RM5575 (TGW), RM1132 (YLD), RM168 (YLD), RM551 (YLD), RM5711 (YLD) and RM204 (YLD). Thirty QTLs were identified as novel based earlier information. At least one QTL was found to be novel for each of 11 traits. Number of novel QTLs found to be 1, 2, 3, 4, 3, 4, 1,2, 4, 1 and 5, respectively for DFF, PH, TLL, PL, FLL, FLW, FG, TG, TGW, SLBR and YLD traits (Table 5, 6).

The association of traits with markers could be confirmed in the 2D plot and Q-Q Plot (Fig 8, S2 Fig, Table 5, Table 6). The QQ-plot showed a similar distribution of marker-trait association for 11 traits (S2 Fig). The GLM Manhattan plot shows 26 SSR markers associated with grain yield at p-value at 0.05 (S3 Fig). However, seven SSRs were associated with grain yield-related traits in MLM Manhattan plot was plotted at p-value 0.005 (S4 Fig). The lowest p-value 8.73E-04 was found with marker RM5709 for plant height trait, followed by 0.00158 (Thousand-grain weight) with RM263 and 0.00266 (Flag leaf width) with RM5709 (Table 5).

### In-Silico Study for Marker Co-localization

The present study has used the computer and web-based servers’ big data to confirm the association of co-localizing genes and QTLs linked with yield-related traits of rice. Twenty SSRs were identified that co-localized with grain yield and related traits (Table 6). Using in-silico approach, it was found that 10 out of 20 co-localized SSR markers were in agreement with previous reports. These 10 co-localized SSRs viz., RM154, RM5709, RM5575, RM20285, RM204, RM3827, RM5711, RM447, RM22899 and RM171 were found to be very important because of their association with grain yield-related traits. RM5709 found to be highest co-localized on 4^th^ chromosome (associated with 9 yield-related traits viz., DFF, PH, PL, FLL, FLW, FG, TG, YLD and SLBR) followed by RM297 (associated with PL, FLL, FLW, TGW and YLD) and RM5575 (associated with PL, FLW, TGW and YLD) (Table 6).

## Discussion

### Phenotypic Variance

Improving rice yield potential is one of the primary breeding objectives in many countries for several decades [5]. In 1960s and 1980s, the green revolution was initiated with the development of semi-dwarf High Yielding Varieties (HYVs) like IR 8 and IR 36 [2,9,55–58]. The main objective of the green revolution was to fulfil and achieve self-sufficiency in food requirement, which helped the developing countries around the world especially in South Asia. It was realized in rice due to development of semi-dwarf, lodging resistant and fertilizer responsive high yielding varieties. It led to stability in rice production and mitigating the hunger of growing population. It was accomplished in mid-sixties with the development of miracle variety IR8. Since then, a stagnant in yield potential of semi-dwarf *indica* inbred varieties was noticed in *indica* inbreds, which needs to be cracked [55].

New Plant Types (NPTs) was a potential approach for breaking the yield ceiling. The initial effort on NPT was made by IRRI scientists to develop 2^nd^ generation NPT genotypes accumulating favorable alleles from tropical *japonicas* and popular *indica* for yield-related traits with multi-environment testing [5, 55]. The main idea behind NPT development was to develop a plant type endowed with combination of unique traits that would help for efficient photosynthesis and partitiining for very high grain yield in irrigated ecology. In this process, favorable *tropical japonica* genes were accumulated in *indica* background in second-generation NPT lines. IRRI scientists identified highly potential genotypes, i.e., IR 72967-12-2-3 which reportedly produce 10.16 ton/ha [9]. Our main target area for breaking yield ceiling was eastern zone of India, which has some climatic constraints particularly low light due cloudy weather in wet season. In current study, the advance generation 2nd generation NPTs were phenotypically screened for high grain yield and associated traits in four seasons at. NRRI, Cuttack, India. Phenotyping is the most crucial step for crop improvement. Identification of suitable transgressive segregants for the specific quantitative trait in any crops is a challenging task for the breeders. At the outset, a set of such elite materials of NPTs was chosen (advance generation segregating materials) as initial materials. Trait specific selection and evaluation of these materials were subsequently led to identification 48 NPTs with variable grain yield were subjected to multi-environment testing. In this study, potential 20 NPTs were identified with an average yield in the range of 5.45 to 8.8 tons/ha. These genotypes could either be utilised directly or prospective parents based on *per-se* yield.

PCA Bi-plot analysis showed association with major yield-related traits (Fig 2B, S5 Table). The PC1 and PC 2 explained 45.67 % and 12.18 % of the total variance, respectively. Similar variability were also reported for PC1 and PC2 viz. 35.2% and 14.4%, respectively [59]. The distribution pattern of traits clearly differentiates the genotypes and relative importance of traits, which influence the grain yield (YLD). The positive relation was observed among genotypes in the first quadrant showed the importance of traits viz., PL, FLL, FLW, TL, YLD, SLBR, and TGW particularly in NPTs. The genotypes are associated with respective traits in a 1^st^ quadrant could be responsible for better average grain yield. The first quadrant consists of important traits and the genotypes endowed with those traits predominantly, hence could be selected as donor parents for specific traits in NPTs. Similarly, traits viz., TG, FG, DFF, and PH were predominant for the genotypes in quadrant IV (S5 Table), and it could be selected as a donor based on a number of fertile spikelets and effective tiller number.

The present study reported that dominant phenotypic traits such as PL, TL, FLL, FG and TG, had a positive correlation with yield. However, the more focused selection should be done for those traits (PH, FLW, TGW and SLBR) that are showing weak correlation with grain yield, because of environmental effects (Table 1, S3 Table). The best genotypes are assigned for grain yield based on phenotypes could be N-129, N-8, and R-255 (Fig 1A and 1B, S2 Table, S3 Table). The dominant specific traits and genotypes for high grain yield could be selected for designing effective breeding strategy. This would be helpful to the breeder for the proper choice of a parent/donors for bi-parental/multi-parental mating vis-à-vis during the process of selection in segregating generations. Therefore, present study reports phenotyping followed by different statistical analysis which suggests trait-specific genotypes for prospective parents in the hybridization programs for breeding super rice [23, 59]

The Heatmap shows a data matrix in the form of the colour pattern due to the numeric differences in multivariate data. In ClustVis, hierarchical clustering can be optionally applied on specific traits those were linked with particular genotypes and observations [23]. The Heatmap analysis showed the order of colour merging with the specific traits that are playing a vital role in the association for targeted trait. The different colouring patterns were linked with respective traits starting from deep green to red (Fig 3). Apart from green, the white and light colour traits indicated relatively poor association with yield-related traits viz., PH, DFF and SLBR traits. As the colour moves along the colour chart from green to red, the association with yield improves with grain yield that means the red colour especially was strongly associated. In this context, the order of association based on the colour intensity starts with FG followed by TG, TGW, FLL, TL, PL and FLW. The respective genotypes have been depicted with their strong associated traits colours. The dense colour which gives ideas for a strong character. So that breeder can easily choose donor parents who are actively associated with specific traits [23] (Fig 3).

### Allelic and Genetic Diversity

The utility of SSR markers for population structure, diversity and association mapping depends on the quality of information they provide. The allelic and genetic diversity helps the breeder to understand genetic constitution of germplasm makeup and target donor selection for designing effective future breeding strategy. The 66 out of 85 SSR markers (77.64%) showed polymorphism, which amplified 154 alleles. Similarly, Anandan and his team reported the 39 polymorphic SSRs which amlified 128 alleles [60]. The average PIC value in this study was found to be 0.70, which was similar to previous reports [23, 59, 61–64]. The lower rate of the average PIC was reported in association studies by several workers {0. 31[65]; 0.47 [66, 67]}. The PIC value showed a positive correlation with the total number of alleles (S6 Table, S7 Table). Similar findings were reported by previous researchers [43, 61]. Moderate levels of genetic diversity (i.e. 0.39) was observed among 60 genotypes used in the present association study. Similarly, Cui et al. (2013) detected an average diversity of 0.34 in 347 genotypes used for association mapping in cold tolerance at the booting stage [65]. However, a higher rate of average genetic diversity was reported by others (0.69: Zhao et al., 2013; 0.52: Nachimuthu et al., 2015; 0.76: Edzesi et al., 2016) [16,67,68].

### Population STRUCTURE

The population structure analysis helps to understand and differentiate the types of population groups exists in a set of genotypes. The population structure based on Bayesian clustering model has been most frequently used to correct spurious associations. The delta K value was measured by ad-hoc and based on the relative rate of change in likelihood LnP (D). The Delta k =2 was set to get a higher likelihood optimal number of LnP (D) among groups.

The 60 genotypes were differentiated into four sub-populations at K=4, i.e., POP1, POP2, POP3, and POP4. Similar sample sizes were used by several researchers in association analysis in rice [69, 70] and alfalfa [71]. The UPGMA cluster analysis grouped 60 genotypes into two major groups at 54 % of genetic similarity. The Nei’s pairwise genetic distance showed three types of populations, i.e. POP1, POP2, and one admixture population. Similarly, at K=2, STRUCTURE analysis could differentiate entire populations into two subpopulations (Fig 7). Genotypes in these populations along with high mean values could be utilized as potential parents for transgressive segregants with high yield and yield attributing traits towards breaking yield ceiling

The mining through the Power Marker into details individual groups, the first population (POP1) contained hardcore NPTs, which was distinctly different from *indica (Ind), temperate japonica (Temp.)* and *tropical japonica (Trop.)* (2^nd^ population, POP2, which also includes few NPTs along with *Ind*, *Temp.* and *Trop*.). However, all NPTs contain the genomes of *indica* as well as *temperate* and *tropical japonicas.* Therefore, the population has one admixture group, which lies in between the two classes comprising the characters of both populations. Therefore, a targeted hybridization between consciously selected parents of these two distant groups might result in transgressive segregants with super rice traits for future yield enhancement. At K=4, the population was clustered into four groups viz., *Ind (1st)*, *Trop (2nd)*, *Temp (3rd)* and *NPTs (4th)* according to their genotypic and evolutionary significance. However, this study suggested that popular varieties clustered together according to their ecology, morphology and inter-varietal hybrid fertility of rice varieties in *indica* and *japonica* [59, 72]. Here, almost all the NPTs were grouped separately, except one i.e., N-129. Moreover, the genotypes in the 4^th^ group comprised the genomes from *indica*, and *japonica* and supposed to have a relationship with the first three clusters. The population cluster 1, 2 and 3 were purer and divergent, but in the 4^th^cluster, genotypes were intermediates of first three clusters. This could help breeders in devising necessary breeding strategy and choosing parents for yield improvement.

### QTL Association

QTL association has been widely used for the identification and mapping of QTLs for various traits such as tolerance to biotic and abiotic stresses, quality and grain yield in different crops [4,6,13,42,49,73]. This study also targets findings new QTLs, alleles, and genes [74] and validate the previously reported QTLs. The present association study was conducted on 60 diverse genotypes panel and 85 SSR markers The present association study focused on identification QTLs associated with yield and related traits in relatively small population and with limited markers [75]. Therefore, our study analogous with previous reports with a small, focused group of genotypes and limited marker pairs combination [12,13,60,67,69,75–77].

In association studies, both GLM and MLM models are used. Population stratification and cryptic relationships are two common reasons for the inflation of false-positive association [39]. GLM model used to have more false positive association as compared to MLM model analysis [20,42,71,78,79]. It does not consider to influence the population structure and kinship [71, 80]. MLM model has higher accuracy and a smaller number of spurious marker-trait association with genotypes as compared to GLM model. This model is having a powerful algorithm, which systematically increases power, improves calibration and reduce computational cost to structured populations generally used for SNPs in GWAS [46, 79]. The MLM model integrates structure and kinship matrix (Q+K) which supposedly corrects the false-positive error to tune of 62.5%. Hence, MLM model has been popularly used in several cases for marker-traits association [12,19,44,46,72, 81–85].

In association mapping, mixed model (Q+K) showed a significant improvement in goodness of fit and reducing spurious associations. The K and Q matrix corrected the association between eleven phenotypic traits with markers [44, 71] at permutation value is 1000 at p<0.005 for GLM and MLM for the level of significance. In association mapping, p-value plays an essential role because it controls over the level of false-positive association between traits and markers. It means that if the p-value is minimized, there is less chance of a false positive association of markers with respective traits and vice-versa [46]. The value of p is in agreement with previous reports [71,72,79,86]. However, some researchers reported their results at p<0.05-0.01 value, which is much higher compared to our study, where the number of markers is appreciably high [44, 71].

GLM identified 30 SSRs which shows 70 associations for grain yield *per se* and other 10 yield-related traits based on the four-season mean data (mean value of 4 seasons) (Fig 8). It was found that 11 commons SSRs were found between GLM and MLM model and have a positive association with yield-related traits based on four-season mean data (Table 5). Twenty-tow SSR markers have been reported previously and these markers have a positive association with QTL regions based on mean data. There were 15 SSRs linked with different QTLs responsible for *per se* grain yield in four different seasons. Out of them, five SSR loci were in aggrement to the previous studies. Previous reports indicated that the markers RM154, RM551, RM5711, RM234 and RM19 were associated with grain yield QTLs, (Table 5) viz., *qYLD2.1* & *qts1*, *qYLD-4* & *qPL-4.1, qYLD-7.1*, and *qYLD-12.1, qSpn-12.1*, respectively [12,25,26,30,59,87]. Similarly, the marker RM5709 has been well documented in the www.gramene.org database for its association with grain yield QTL. Five SSR markers, RM6100, RM1132, RM5575, RM20285 and RM26499 are linked with novel QTLs, *qYLD10.1, qYLDa7.1 qYLD-5.1, qYLD6.1* and *qYLD11.1* responsible for imparting high grain yield. In case of tiller number, three out of four SSR markers RM26499, RM3276 and RM3827 were found to be associated with the novel QTLs, i.e. *qTL-11.1, qTL-4.1* and qTL*-6.1*, respectively. For panicle length(PL), previous researchers have reported the association of RM5709 with *qPL-4.1*; *qSPP-4.1*, RM204 with *qPL-6.1* [3], RM234 with *qPPL-7.2* [30]; RM447 with *qPL-8* [21, 88] and RM17 with *qPL-12b*. Four out of 13 SSRs viz., RM297, RM5575, RM5711, and RM22899 were found to be associated with four novel QTLs, *qPL-1.1, qPL-5.1* and *qPL-8.1*, respectively for panicle length (PL). However, Marathi et al. (2012) reported that RM5709 marker was linked with *qSPP-4.1*, indicating the pleiotropic effect of *qPL4.1* on panicle length. The present study reported that a total of 14 SSRs were associated with FLW out of them 3 SSRs RM297, RM5709, RM1132) were reported previously [28,29,31,67,68]. Four SSRs, RM447, RM297, RM5709 and RM285 have been identified for flag leaf length (FLL) in QTL regions of *qFLL8.1*, *qFLL-1.1* [29], *qFLL-4.1a* and *qFLL9.1*, respectively. Out of the above, two markers, RM297 and RM5709 were also associated with QTLs for flag leaf width(FLW), *qFLW-1.1* and *qFLW4.1*, respectivelly due to pleiotropic effects [28, 29]. Similar reports corroborate the present finding of marker-trait association for RM1132 apart from three other reported markers [31]. It is suggested that one locus may be involved in conferring multiple traits, which may be the result of the gene to gene interactions and pleiotropic effect. Three markers QTL associations viz., RM154∼ *qFLW2.1*, RM168 ∼*qFLW3.1*, and RM470 ∼ *qFLW4.1* were identified for the flag leaf width (FLW) (http://www.gramene.org/). Four novel QTLs, *qFLW-6.2, qFLW-5.1, qFLW-8.1* and *qFLW-7.1* were identified for controlling FLW trait in our study.

The RM5709 was reported to be linked with QTL, *qPHT4-a* for plant height [28]. Number of fertile grains is considered as important trait, because of its link with grain yield. Same SSR marker (RM5709) was also found to be associated with *qFGP4.1* [28]. Thousand-grain weight (TGW) is another crucial trait supposedly linked to yield. Our study identified seven QTLs viz., *qTGW-a1.1, qTGW-a9.1, qTGW-a9.2, qTGW-a10.1, qTGW2.1 qTGW-5.1* and *qTGW-8.1* associated with SSRs viz., RM297, RM201, RM219, RM171, RM263, RM5575 and RM22899, respectively. These QTLs might be highly useful in the rice breeding programs. Four QTLs were found to be novel i.e., *qTGW-a1.1, qTGW-a9.2, qTGW-5.1* and *qTGW-8.1*, which could be emphasized because of their better grain filling and boldness leading to higher grain yield. Marri et al. (2005) reported that RM263 was found to be linked with *qTGW-2.1, qGn2.1*, and *qYLD-2.1*, indicating the possibility of pleiotropic effects. *qGn2.1* was also reported earlier [49],

Twenty SSRs were having association with more than one traits and have been reported in the gramene-database (http://www.gramene.org/) by earlier studies (Table 6) [28,29,49,89]. These were significantly associated with yield controlling complex traits viz., PH, DFF, SLBR, TL, PL, FLL, FLW, FG, TG, TGW and YLD and supposed to have played a significant role in yield enhancement (Table 5, Table 6, Fig 8). Similar reports by Zhang and team (2014) revealed the pleiotropic effect of gene *LSCHL-4*, in influencing increment of leaf chlorophyll, enlargement of flag leaf size, higher panicle branch and grains per panicles [31]. The previous report suggests that specific marker association with more than one traits might be either due to the linkage of genes or pleiotropic effects of a single locus [90–92]. However, variation in population structure, QTL detection methods and environmental conditions restrict our choices to compare the newly identified QTLs with the already reported QTLs. Therefore, we do have a need for further study on these potential trait-specific genotypes. It would lead to design effective breeding strategy for introgression of high grain yielding traits associated QTLs into popular rice varieties for obtaining super rice targeting to overcome yield ceiling.

The in-silico study of co-localization of SSR markers with respective traits would be helpful to the breeders to confirm the association of trait-specific SSRs. The present study has reported 10 SSR markers, which are reported to be associated with grain yield-related traits and found with gene IDs (Table 6). Most of the SSRs were co-localized with more than two traits. The highest co-localization was identified in RM5709 linked with nine traits followed by RM297 co-localized with five traits. The earlier study reported that RM5709 was associated with gene ID Os04g0619500 and it express in ovarian tumour, which produce domain-containing protein [52,53,93]. Similarly, RM297 was associated with five traits viz., PL, FLL, FLW, TGW, YLD, while RM5575 and RM204 both are associated with four traits (Table 6). The previous study reports the marker RM5575 linked with gene Os05g0474600, which produces, similar to Aldose reductase-related protein [52,53,93]. The RM204 was associated with four traits viz., PL, FLW, YLD and SLBR. International rice genome sequencing project IRGSP-1 (2005) reported that [51], RM204 was linked with gene Os06g0162800, which helps in mads-box transcription factor, and controls spikelet morphogenesis along with regulation of floral meristem [51–54,93]. Similarly, RM22899 was found to be associated with three traits viz., PL, FLW and TGW, and also linked with gene Os08g0331900, and it induces the expression of similar to Leucine Rich Repeat family protein [53, 93]. The current In-silco study helps us for confirmation of 10 colocalized SSRs that were associated with grain yield-related traits. This marker could be useful in marker-assisted back cross-breeding program to produce next-generation super rice

## Conclusion

Breaking yield ceiling in rice warrants conscious selection of parents. Sufficient variability was available in NPT genotypes, because of the genetic distance of parents using tropical *japonicas* and *indicas*, leading to fixation of distinct lines. In the present study, microsatellite markers were used for association studies for grain yield and ten yield-related traits. Few highly-potential genotypes with high yield along with variation in morpho-physiological traits were identified after conducting the trial in four consecutive years. Wide variations were found in all the traits, which would be helpful for the identification of genotypes required for bi-parental/multi-parental crosses in developing super rice genotypes with higher grain yield. The STRUCTURE and tree diagrams were helpful in the classification of populations into distinct clusters vis-à-vis uniqueness among them and helped in the identification of a diverse gene pool for necessary parental selection for targeted transgressive segregants. The in-silico study reported twenty SSR markers, those were associated with more than one trait. Nineteen, SSR markers were found to be associated with different traits in more than one season. Four SSR, RM5709, RM5575, RM20285 and RM234 were found to be associated with PL, PH, SLBR and YLD and common for four seasons. More than 25% phenotypic variance was explained by QTLs bracketing RM5709 for each of nine traits, DFF, PH, PL, FLL, FLW, FG, TG, SLBR and YLD. RM5709 would be useful for transfer above nine traits into popular rice varieties. The present study reported that 16 SSRs were linked with 11 yield traits and were found to be associated with 30 novel QTLs. Some of the QTLs are very important viz., TL (*qTL-6.1*), PL (*qPL-1.1, qPL-5.1, qPL-7.1, qPL-8.1*), FLL (*qFLL-9.1*), FLW (*qFLW-5.1, qFLW-8.1*), TGW (*qTG-2.2, qTGW-5.1, qTGW-8.1*), SLBR (*qSlb-3*) and YLD (*qYLD-5.1, qYLD-6.1a, qYLD-7.1, qYLD-11.1*) because of their association with important yield contributing traits or yield *per se* for multiple seasons. Hence, these could be pyramided in elite background for realization of higher yield and breaking yield ceiling. The study would be immensely helpful for selecting target donors with requisite traits for designing effective future breeding strategy for super rice.

## Abbreviation

NPT-New plant type; GLM-General Linear Model; MLM-Mixed Linear Model; MS-Mean sum of square; Fst-F-statistics, subpopulations within the total population, Fis-F-statistics; individuals within subpopulations; DFF-Days to fifty percent flowering; PH-Plant height; TL-Tiller number; PL-Panicle length; FLL-Flag leaf length; FLW-Flag leaf width; FG-Fertile grains; TG-Total number of grains; TGW-1000 grains weight; YLD-grain yield t/ha; SLBR-Seed length-breadth ratio; FDR-False Discovery Rate.

## Author’s Contributions

**SKD, RD and LB-**conceptualization and designing of experiments; **RD, SR-**Performed the research, data analysis; **RD-** Performed the research on Interpretation, writing manuscript; **RD, SM**-Performed the research on genotyping; **SKYB, BP, MM, KC, AA-**Performed research on phenotyping experiments and data collection; **SKD, ONS, PS, LB-**Performed correction of manuscripts; **SKD, LB and KKS-** Conducted the field Experiment and supervision of experimental works.

## Acknowledgements

We thank the IRRI for providing the NPT seeds and Director, ICAR-National Rice Research Institute (NRRI), India, for providing all laboratory and field facilities.

## Supporting Information

**S1 Fig.**
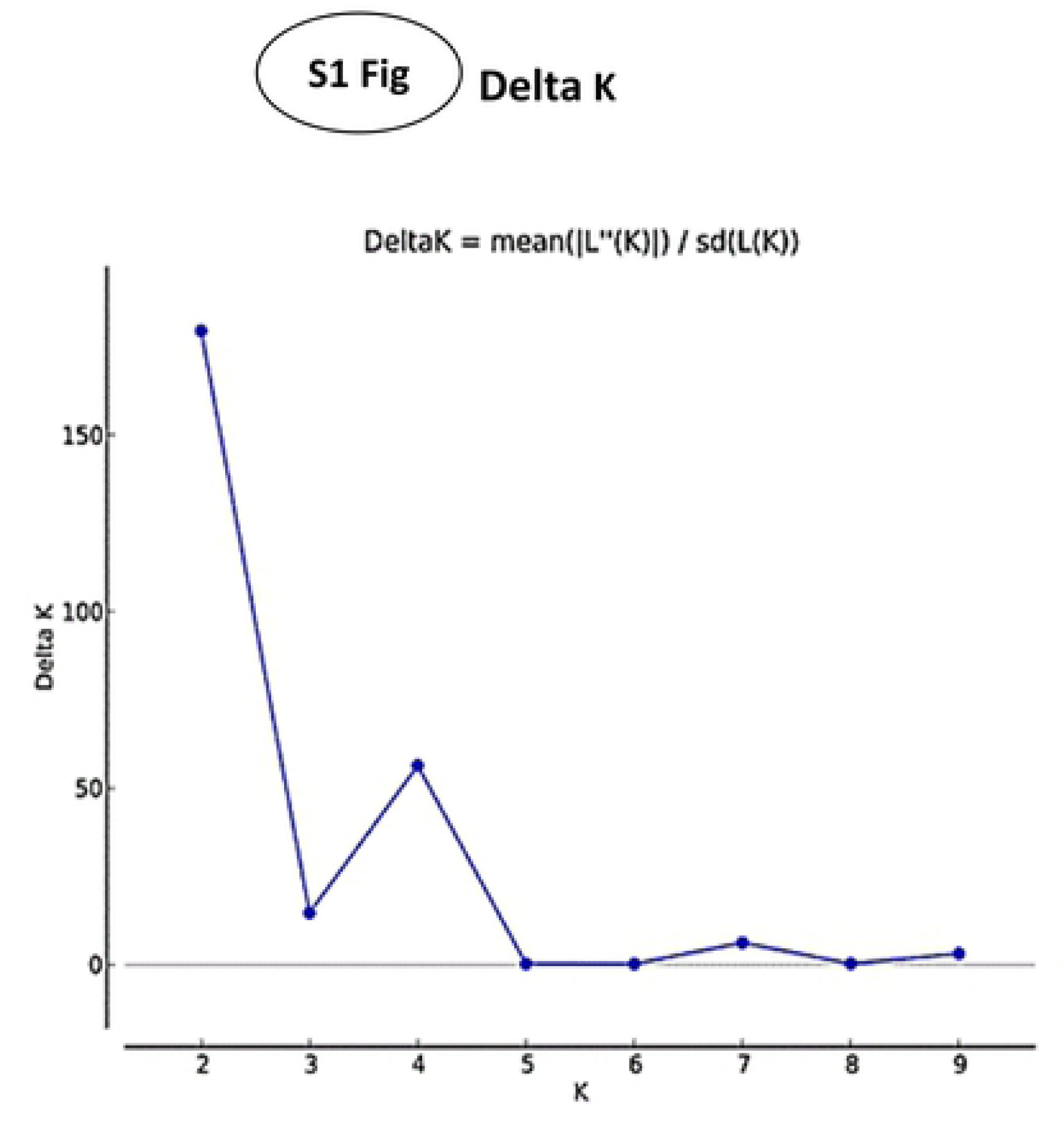
The true value of K was determined by STRUCTURE harvester in K-2 Plot of change in the likelihood of the data, L (K), at values of K from 1 to 10 K=2.

**S2 Fig.**
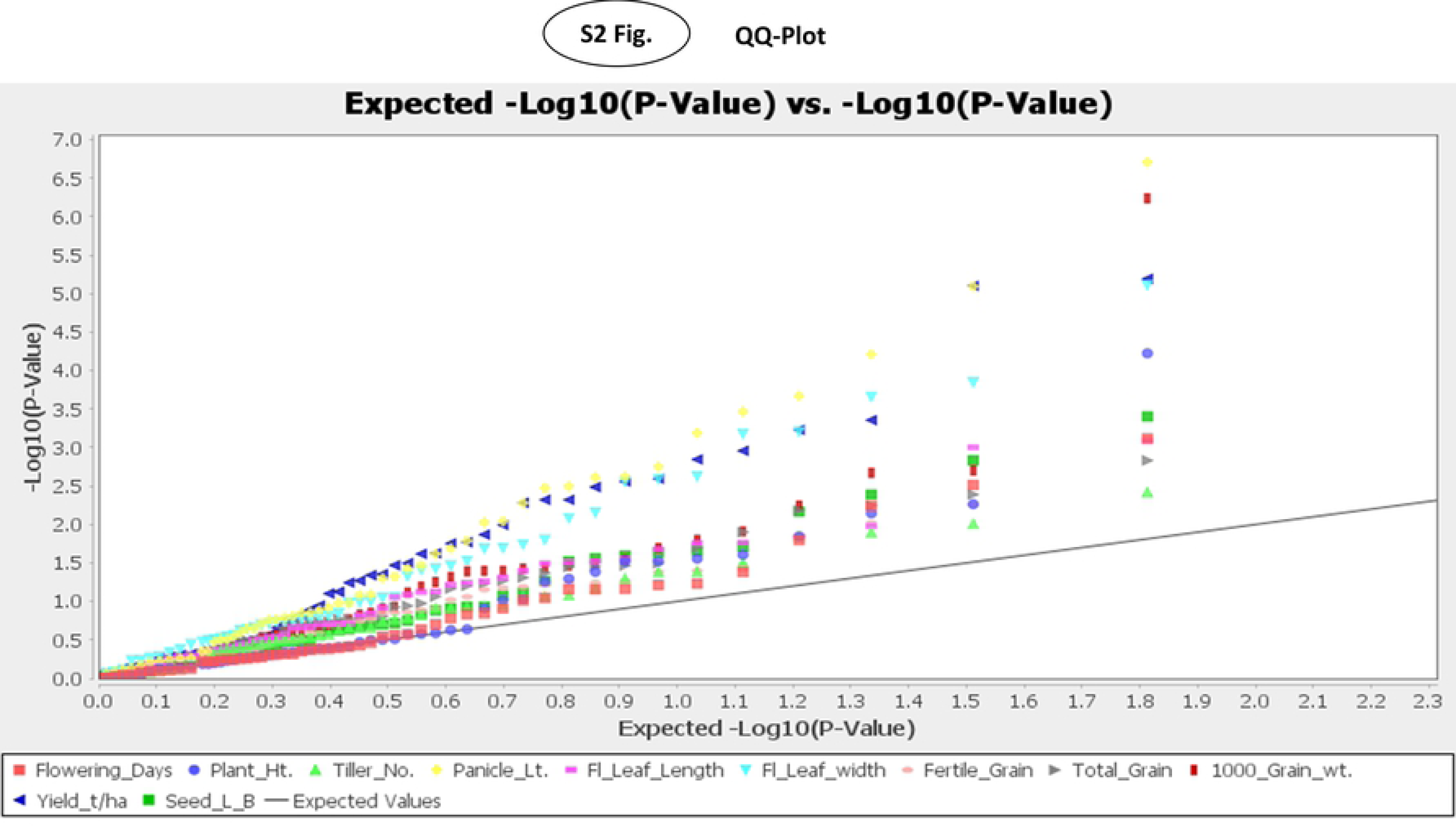
QQ-plot showing distribution of marker-trait association for 11 traits.

**S3 Fig.**
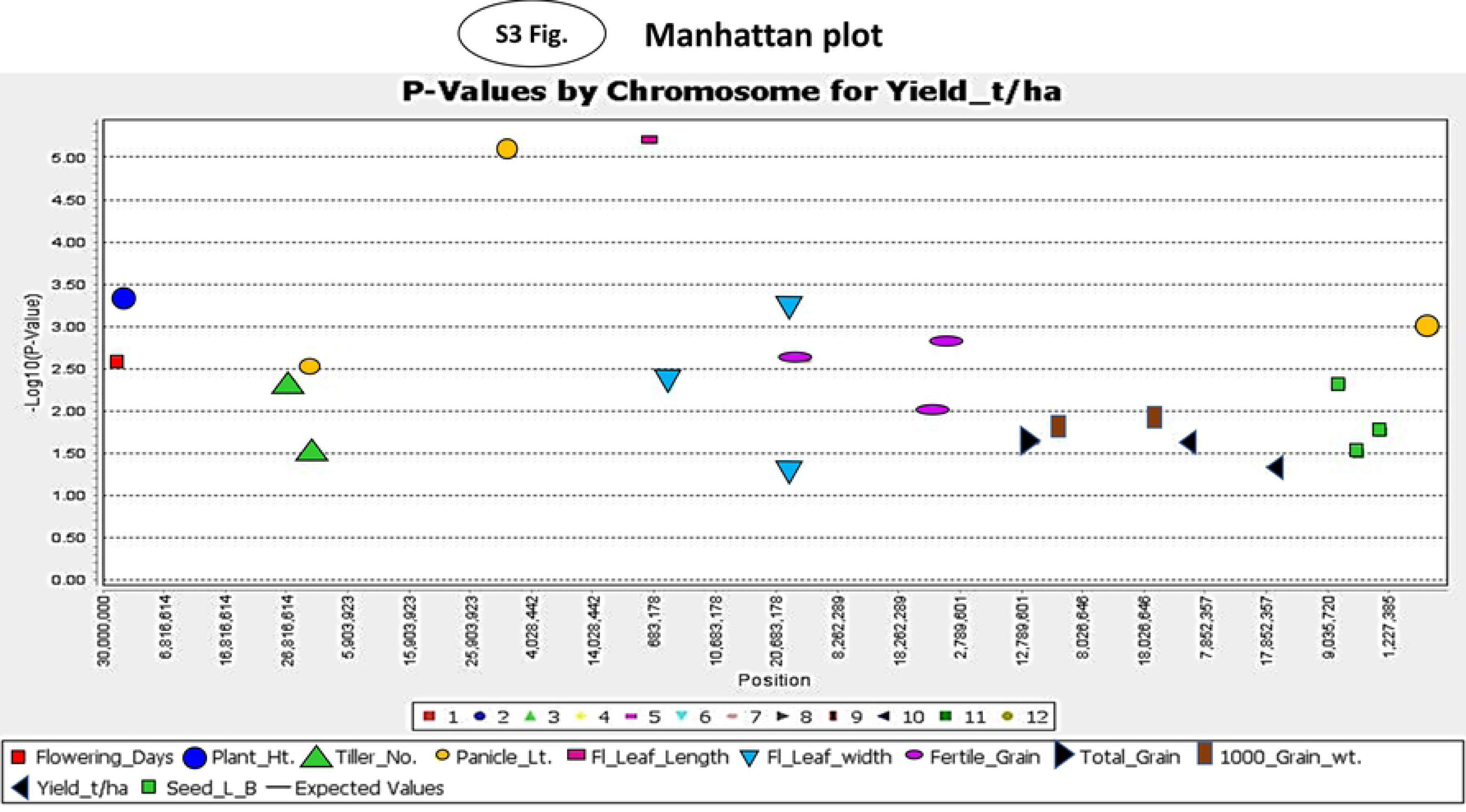
GLM Manhattan plot showing markers associated with grain yield using significant p-value at 0.05.

**S4 Fig.**
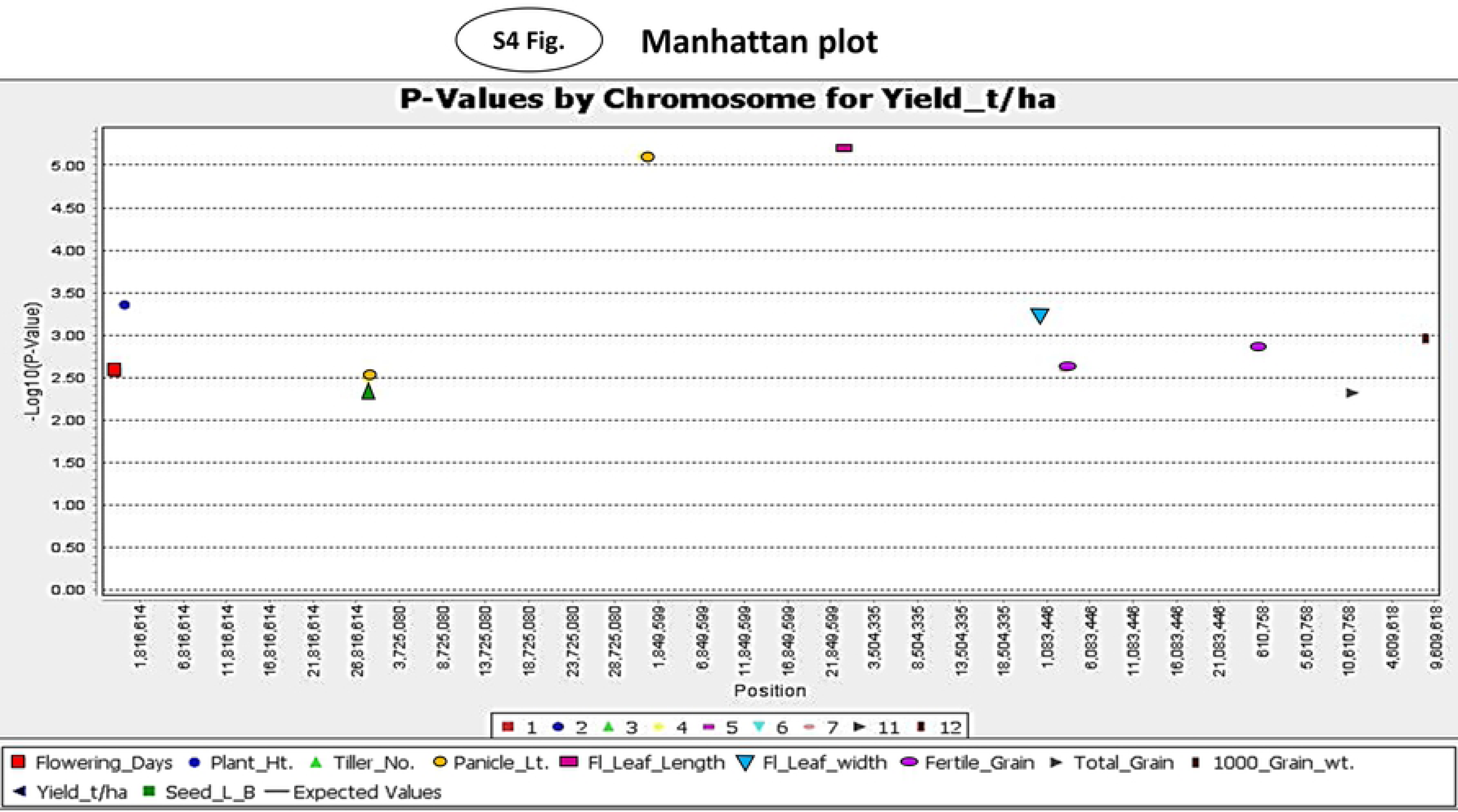
MLM Manhattan plot showing markers associated with grain yield using significant p-value at 0.005.

**S1 Table. List of 48 New plant types (NPT) and 12 popular DUS reference genotypes used in association mapping analysis.**

**S2 Table. Grain Yield performance of 20 best varieties and standard checks under irrigated condition.**

**S3 Table. Correlation matrix of grain yield and their association with 10 yield-related traits**

**S4 Table. Calculation of Standardized coefficients**

**S5 Table. Distribution pattern of genotypes in Principal Component Analysis (PCA) and Biplot by using morphological-physiological data**

**S6 Table. Molecular diversity among 60 rice genotypes based on the alleles amplified by 66 polymorphic SSR markers**

**S7 Table. Correlation between PIC and different types of alleles**

